# Two cowpea Rubisco activase isoforms for crop thermotolerance

**DOI:** 10.1101/2025.01.31.635870

**Authors:** Armida Gjindali, Rhiannon Page, Catherine J. Ashton, Ingrid Robertson, Mike T. Page, Duncan Bloemers, Peter D. Gould, Dawn Worrall, Douglas J. Orr, Elizabete Carmo-Silva

## Abstract

- A vital crop for sub-Saharan Africa, cowpea productivity is threatened by climate change, including increased heatwave intensity, duration, and frequency. Rubisco activase (Rca) is a key molecular chaperone that maintains Rubisco activity and due to its thermal sensitivity is a key target for improving crop climate resilience.
- We identified and characterised four Rca isoforms in *Vigna unguiculata* (L.) Walp. cv. IT97K-499-35 *in vitro*: Rca1β, Rca8α, Rca10α and Rca10β. Cowpea leaf and plant growth traits were also investigated during a 5-day +10 °C heatwave that included assessment of Rubisco and Rca activity in leaf extracts, and relative changes in the abundance of the four Rca isoforms.
- Cowpea Rca10α and Rca10β had higher thermal maxima, broader thermal optima and higher rates of ATP hydrolysis and Rubisco reactivation *in vitro*. In the absence of water deficit, the heatwave caused only mild effects, including increases in leaf temperature and expression of *Rca10*, small decreases in Rubisco activity and activation state, and unaltered temperature response of Rubisco activation by Rca.
- The superior Rca10α and Rca10β isoforms offer the prospect of enhancing the thermotolerance of cowpea and other crops in anticipation of more extreme future heatwaves.

## Introduction

Changes in global temperatures including heatwave events pose significant threats to global food security (Moore *et al*., 2021), making increased crop resilience against heat stress imperative (Song *et al*., 2023). Regions closer to the equator, particularly sub-Saharan Africa, are the most vulnerable to rising temperatures (Harboure, 2024). Elevated temperatures can lead to decreased photosynthesis and plant growth (Hatfield & Prueger, 2011; Bagati *et al*., 2021; Zahra *et al*., 2023; Becker *et al*., 2023). Cowpea (*Vigna unguiculata* (L.) Walp) is a major source of protein for populations in sub-Saharan Africa and, despite being adapted to a warm climate, it is still vulnerable to heat stress – even a 1 °C increase in night temperature can result in significant yield loss (Warrag & Hall, 1984; Gonçalves *et al*., 2016; Mohammed *et al*., 2024). Therefore, it is imperative to identify physiological and biochemical traits that confer thermotolerance in cowpea to develop heat resilient crops (Barros *et al*., 2021).

Moderate heat stress, defined as temperatures up to 40 °C (Crafts-Brandner & Salvucci, 2000; Kumar *et al*., 2009; Ara *et al*., 2013), has a profound impact on plant photosynthesis, adversely affecting both the electron transport chain and the Calvin-Benson-Bassham (CBB) cycle (Chen *et al*., 2022). Rubisco and chlorophyll leaf content also decline under moderate heat stress (Mathur *et al*., 2014). Chlorophyll biosynthesis enzymes exhibit susceptibility to heat stress; however, the reduction in Rubisco content cannot be attributed to denaturation of Rubisco, as alterations in secondary structure are observed only at temperatures exceeding 45 °C (Li *et al*., 2002; Mathur *et al*., 2014). Increased temperatures enhance Rubisco carboxylation activity; but also lead to reduced Rubisco specificity and increased O_2_ diffusion, both of which result in higher oxygenation over carboxylation and increased photorespiration (Hermida-Carrera *et al*., 2016; Galmés *et al*., 2016). Importantly, photorespiration does not fully account for the decline in carbon assimilation during moderate heat stress, as the Rubisco activation state has been demonstrated to decrease under both photorespiratory and non-photorespiratory conditions during mild heat stress (Feller *et al*., 1998; Crafts-Brandner & Law, 2000; Galmés *et al*., 2016).

Numerous studies have demonstrated that Rubisco inactivation is one of the primary causes of reduced photosynthetic efficiency during moderate heat stress (Feller *et al*., 1998; Crafts-Brandner & Law, 2000; Portis, 2003; Ristic *et al*., 2009; Carmo-Silva *et al*., 2015; Perdomo *et al*., 2017; Wijewardene *et al*., 2021; Scafaro *et al*., 2023). To catalyse carboxylation or oxygenation, Rubisco catalytic sites require binding of CO_2_ and Mg^2+^ as cofactors to form the active carbamylated Rubisco prior to binding the substrate ribulose 1,5-biphosphate (RuBP). Rubisco can become inactivated by the binding of inhibitory sugar phosphates to carbamylated catalytic sites or by misfire products that cannot dissociate and maintain catalytic sites in their closed, inactive conformation (Orr *et al*., 2023). Inactivation can also occur when RuBP binds to an uncarbamylated Rubisco catalytic site. Inhibited Rubisco requires its molecular chaperone Rubisco activase (Rca) to restore its activity (Carmo-Silva & Salvucci, 2013). Rca is a nuclear encoded protein that belongs to the AAA+ ATPases superfamily (Hazra *et al*., 2015) and, in its active multimeric form, exhibits two activities: ATP hydrolysis and Rubisco reactivation (Bhat *et al*., 2017; Serban *et al*., 2018). Rca docks on Rubisco’s large subunit and, using ATP, performs conformational remodelling to release the inhibitory compound, thus modulating Rubisco activity by facilitating the dissociation of bound inhibitors. Elevated temperatures increase the frequency of Rubisco inactivation due to higher concentrations of some inhibitory substrates (Salvucci & Crafts-Brandner, 2004, p. 20). Furthermore, although Rubisco is a relatively thermostable enzyme, Rca is thermolabile and even a moderate increase in temperature results in loss of its activity (Crafts-Brandner & Salvucci, 2000; Salvucci *et al*., 2001).

Rca isoforms within a single species vary in their regulatory properties (Zhang & Portis, 1999; Carmo-Silva & Salvucci, 2013; Perdomo *et al*., 2019; Scafaro *et al*., 2019b). Many species express two types of Rca isoforms: longer α isoforms (43-47 kDa) and shorter β isoforms (41-43 kDa) that can be encoded either by individual genes or the same gene via alternative splicing (Carmo-Silva *et al*., 2015). Rca α isoforms contain a C-terminal extension with two cysteine residues shown to form a disulfide bond which is regulated by thioredoxin-f in *Arabidopsis* (Zhang & Portis, 1999). This regulatory mechanism links Rubisco activity with alterations in the redox potential induced by environmental stimuli, such as high light (Zhang *et al*., 2002; Perdomo *et al*., 2021). The redox regulation is species-specific, with Solanaceae species such as tobacco (*Nicotiana tabacum* L.) and tomato (*Solanum lycopersicum* L.) containing only the β Rca isoform that lacks the cysteine residues and the redox regulation of the enzyme (Carmo-Silva & Salvucci, 2013). In species containing only redox insensitive Rca, light regulation appears to be mediated by changes in the ADP/ATP ratio of the chloroplast (Carmo-Silva & Salvucci, 2013).

In addition to variation in the number of genes and the isoforms they produce, the expression and relative abundance of Rca isoforms vary among species and as a response to temperature (Law & Crafts-Brandner, 2001; Wang *et al*., 2010; Degen *et al*., 2020; Perdomo *et al*., 2021). In rice (*Oryza sativa* L.) two Rca isoforms are produced via alternative splicing and while Rca β is more abundant under optimal steady state conditions (Wang *et al*., 2010), heat stress induces upregulation of Rca α (Scafaro *et al*., 2016). In maize (*Zea mays* L.) the two isoforms derive from two genes *ZmRcaα* and *ZmRcaβ* encoding an α (43 kDa) and a β (41 kDa) isoform respectively (Yin *et al*., 2014). Under optimal temperature conditions, the ZmRca β protein isoform is expressed in higher levels than ZmRca α and both exhibit comparable diurnal variation in expression levels (Ristic *et al*., 2009). Jiménez *et al*. reported an increase in ZmRca α in seedlings exposed to higher temperatures, however in more mature heat-treated plants this effect was not consistent (Ristic *et al*., 2009; Stainbrook *et al*., 2024). In wheat (*Triticum aestivum* L.) there are two *Rca* genes, one of which produces a β isoform while the other (*TaRca2*) is alternatively spliced and produces both an α and a β isoform (Nagarajan & Gill, 2018). The Rca α/β ratio changes when wheat is exposed to heat stress, suggesting that the relative abundance of the different Rca isoforms is linked with the physiological response to heat stress (Warrag & Hall, 1984; Crafts-Brandner & Salvucci, 2000). By contrast, in *Arabidopsis* the Rca isoform ratio is not impacted by heat, with the α and β isoforms originating from alternative splicing continuing to be expressed at a 1:1 ratio (Zhang & Portis, 1999). These species-specific responses of Rca and Rubisco regulation to heat stress illustrate the need to investigate this system directly in the crop of interest.

While the exact mechanism of Rca deactivation at high temperatures is still not fully understood, it is evident that the optimum temperature range for Rca is species-specific and in tune with the photosynthetic optimum (Scafaro *et al*., 2016). Therefore, the temperature at which Rca loses its activity depends on environmental acclimation; i.e., plants that grow at high temperatures, such as cotton and agave, maintain Rca activity at higher temperatures (Salvucci & Crafts-Brandner, 2004; Carmo-Silva & Salvucci, 2011; Scafaro *et al*., 2016; Shivhare & Mueller-Cajar, 2017). Rca thermotolerance also differs between closely related species and among the native Rca isoforms within a plant species. The wild rice species *Oryza australiensis* Domin and *Oryza meridionalis* Ng. contain thermostable Rca isoforms that enable sustained high Rubisco activity and photosynthetic rate even at 45 °C (Scafaro *et al*., 2016, 2018). In rice, spinach and cotton, the α isoform exhibits higher thermostability whereas in wheat, it is one of the two shorter β (TaRca1β) isoforms that is most thermostable (Kurek *et al*., 2007; Wang *et al*., 2010; Keown & Pearce, 2014; Degen *et al*., 2020). Increasing the thermostability of Rca is considered one of the most promising strategies for increasing Rubisco activity and enhancing photosynthesis under elevated temperature (Carmo-Silva *et al*., 2015; Wijewardene *et al*., 2021).

In this study we identified four Rca isoforms in cowpea encoded by three genes: *Rca1*, *Rca8* and *Rca10*, with the latter producing two isoforms via alternative splicing. Characterisation of their ATPase and Rubisco reactivation activities across a range of temperatures *in vitro* revealed that the two Rca10 protein isoforms have higher thermal maxima and broader thermal optima for both activities. We then examined the effect of moderate heat stress during a 5-day heatwave at 38 °C (+10°C compared to control) on the expression and abundance of Rca isoforms and subsequent impacts on Rubisco activation in the leaves of young cowpea plants. Heat treatment resulted in decreased Rubisco activation state and upregulation of *Rca8a* and *Rca10*, however *Rca10* levels were consistently the lowest. Despite upregulation of the more thermotolerant isoforms, the temperature profile of Rubisco reactivation by Rca in leaf extracts remained unaltered in heat-treated plants. Leaf temperature increased by 6 °C during the heatwave, reaching 32 °C, which is within the temperature range where Rca activity is maintained above 70% of the maximum (T_opt_) for all isoforms. We therefore conclude that warm-adapted cowpea plants can cope with the mild heat stress induced by a 5-day heatwave at 38 °C in the absence of water deficit, which did not impair Rca function. Overexpression of the most thermotolerant, yet least abundant, Rca isoforms in cowpea will enable adaptation to leaf temperatures resulting from heatwaves that are more intense or accompanied by water deficit, which are likely to occur in sub-Saharan Africa.

## Materials and Methods

### Gene identification

Cowpea *Rca* genes were identified by searching the cowpea genome (Phytozome 12, cowpea genome v1.1) using keywords and known Rca protein sequences from *Arabidopsis thaliana*, *Nicotiana tabacum*, and *Triticum aestivum* (Supporting Information Table S1)(Goodstein *et al*., 2012; Lonardi *et al*., 2019). This identified 3 genes for *Rca* in cowpea: *Vigun01g219300* (*Rca1*), *Vigun08g150700* (*Rca8*), and *Vigun10g051600* (*Rca10*). Alignment with known sequences and key conserved features of canonical Rcas allowed identification of the products of each gene. Transit peptide sequences for each protein were identified by alignment to known Rca sequences (with known transit peptide locations). The analysis was repeated using the updated Phytozome 13 cowpea genome v1.2 and yielded identical Rca isoforms (Supporting Information Fig.S1). Analysis of transcript sequences showed that *Rca1* encodes a 384 amino acid (aa) mature protein consistent with a β isoform. *Rca8* encodes a 419 aa α isoform and a 295 aa isoform that is shorter than a regular β isoform and unlikely to be functional. *Rca10* encodes both a 419 aa α and a 382 aa β isoform by alternative splicing.

The cowpea *HSP20* (*Vigun06g052200*) gene, used as a heat stress marker, was identified as a homolog of the soybean *HSP20* (*Glyma.18g094600*) shown to be upregulated in two soybean cultivars exposed to heat stress (Song *et al*., 2017). The Arabidopsis homolog (*At4G27670*) is also upregulated in heat stress (eFP Browser Stress Series, https://bar.utoronto.ca/)(Kilian *et al*., 2007; Winter *et al*., 2007). The *Glyma.18g094600* coding sequence was downloaded from SoyBase (https://legacy.soybase.org) and used to search the cowpea genome as above (Brown *et al*., 2021).

### Cloning of cowpea Rca and construction of *E. coli* expression vectors

Coding regions for the mature proteins of cowpea Rca1β, Rca8α, Rca10α and Rca10β were codon optimised for *E*. *coli, de novo* synthesised and subcloned into pUC57 (Kan^R^) by Aruru Molecular Ltd. (Dundee, UK). A Golden Gate compatible *E.coli* expression vector was created from the pET His TEV LIC plasmid (a gift from Scott Gradia, Addgene plasmid #29653; http://n2t.net/addgene:29653; RRID:Addgene_29653). Ligation independent cloning was used to add a Golden Gate cassette taken from pICH47742 (part of the MoClo Toolkit that was a gift from Sylvestre Marillonnet, Addgene kit # 1000000044) into pET His TEV LIC, generating the new Golden Gate compatible pET His TEV GG plasmid (Weber *et al*., 2011; Werner *et al*., 2012). Cloning a coding region into this vector allows production of a target protein fused to an N-terminal histidine (His) tag that can be removed with TEV protease. PCR was used to add Golden Gate overhangs to the four optimised *Rca* sequences (see Supporting Information Table S2 for primers used). Golden Gate reactions (final volume 15 µL) were performed with a 2:1 molar ratio of insert:plasmid and the following reagents (final concentration): 1× T4 DNA Ligase buffer, 0.1 mg mL^-1^ BSA, 1 mM ATP, 40 KU/mL T4 DNA Ligase, 2 KU/mL *Bsa* I. All reagents from NEB, except ATP (Promega). Reaction conditions were 30 cycles of [37 °C for 2 min followed by 16 °C for 3 min], 37 °C for 5 min, 80 °C for 10 min, and 16 °C for 1 min. 5 µL of the reaction mix was used to transform *E*. *coli* DH5α cells. Resulting colonies were used to generate plasmid DNA for sequencing with primers T7F and T7R (Source Bioscience) to verify correct sequence including in-frame fusion of the coding region to the His tag.

### Recombinant Rca expression and purification

Expression constructs were used to transform BL21(DE3)pLysS *E. coli* competent cells (Invitrogen). PCR and sequencing were used to confirm the presence of the correct constructs in the BL21(DE3)plysS *E. coli* cells. Several cell lines were tested for each construct to identify the cell strains with the highest expression levels for the target proteins. Expression experiments were then scaled up for protein purification.

Starter cultures (10 mL M9ZB medium, containing kanamycin and chloramphenicol, both at 50 µg/mL) were inoculated from glycerol stocks and grown overnight at 37 °C with 225 rpm orbital shaking. Dynamite medium (containing kanamycin and chloramphenicol, at 50 µg/mL) was seeded with the overnight starter culture (10 mL starter culture in 1 L Dynamite medium). This culture was grown for 3-4 h at 37 °C with shaking at 220 rpm until OD_595_=0.6-0.8 and then IPTG (0.5 mM final concentration) added to initiate protein expression. Cultures were then transferred to 24 °C with 220 rpm shaking for 20 h. Cells were pelleted by centrifugation (5,000 × *g*, 4 °C, 20 min), washed with 40 mL buffer (50 mM HEPES pH 7.2, 300 mM NaCl, 5 mM MgCl_2_), transferred to 50 mL screw capped tubes before finally collecting by centrifugation (5,000 × *g*, 4 °C, 20 min). The pellet was flash frozen in liquid nitrogen and stored at −80 °C.

Cell pellets were thawed on ice and resuspended in Lysis Buffer (50 mM HEPES pH 7.2, 5 mM MgCl_2_, 300 mM NaCl, 50 mM Imidazole, 0.1% Triton X-100 (v/v), 10 µM leupeptin, 1 mM PMSF), using 7 mL buffer for every 1 g of pellet. Cells were lysed on ice using a probe sonicator and 5 cycles of 30 s on 30 s off until the solution was homogeneous. Cell debris was removed by centrifugation (23,000 × *g*, 4 °C, 30 min) and the soluble fraction pushed through a 0.45 µm syringe filter prior to chromatography.

Purification of the Rca protein was achieved using an NGC chromatography system (Bio-Rad, UK), equipped with a nickel-charged IMAC column (1 mL HisTrap Fast Flow, Cytiva) and a desalting column (10 mL Bio-Scale Mini Bio-Gel P6 Desalting Cartridge, Bio-Rad). IMAC and desalting columns were equilibrated with Equilibration Buffer (5 mL: 50 mM HEPES pH 8, 300 mM NaCl, 5 mM MgCl_2_, 50 mM imidazole) and Final Buffer (13 mL: 50 mM HEPES pH 8, 5 mM NaCl, 5% glycerol), respectively. The sample was applied to the IMAC column, followed by Wash Buffer (15 mL: 50 mM HEPES pH 8, 300 mM NaCl, 5 mM MgCl_2_, 125 mM imidazole). Bound protein was eluted using a higher concentration of imidazole (2.1 mL: 50 mM HEPES pH 8, 300 mM NaCl, 5 mM MgCl_2_, 300 mM imidazole) and directly applied to the desalting column. Buffer was exchanged with 15 mL Final Buffer and 1 mL fractions collected. Fractions containing protein were identified using a UV/Vis detector reading at 280 nm. Fractions containing the highest protein concentration (determined using Bradford reagent, BIO-RAD) were pooled, ATP and DTT (2 mM and 5 mM final concentration respectively) added and 50-100 µL aliquots frozen in liquid nitrogen before storage at −80 °C.

### *In vitro* characterization of Rca thermal optima

The rate of ATP hydrolysis by Rca isoforms was assayed using the method described by Chifflet *et al*. (1988), with modifications as described within Degen *et al*. (2020). ATP hydrolysis was measured at a range of temperatures (20-55 °C) using 5-minute assays, with reactions initiated through addition of Rca (final concentration 1 µM Rca monomer) and quenched via addition of SDS. After quenching, all reactions and standards were kept at 4 °C until inorganic phosphate (P_i_) determination. The P_i_ released during ATP hydrolysis was determined by measuring the absorbance of molybdenum blue at 850 nm in a spectrophotometer (SpectroStar Nano, BMG Labtech, UK).

Rubisco reactivation assays were conducted at a range of temperatures (20-45 °C) following the procedures of Barta *et al*. (2011) and Degen *et al*. (2020). Briefly, Rubisco purified as per Amaral *et al*. (2024) was used to prepare inhibited Rubisco complexes (ER) through incubation with RuBP overnight at 4 °C. The two-stage reactivation assay comprises a first stage assay where Rca is added to inhibited Rubisco (ER) at a test temperature (20-45 °C), with aliquots from this assay taken at time intervals to initiate the second stage assay that measures Rubisco activity. This Rubisco activity is a function of the Rca, time allowed for reactivation, and the temperature of the first stage (Supporting Information Fig. S2). With known quantities of Rca and Rubisco, and appropriate controls as detailed within Degen *et al*. (2020), Rca reactivation of Rubisco can be established. Maximum Rubisco activity was determined at every temperature and was comparable among assays with each of the four Rca isoforms.

Independent purifications of each isoform were used as biological replicates. For ATPase activity, three biological replicates were measured for Rca8α and four for the each of the rest of the isoforms. For Rubisco reactivation, four biological replicates were measured for temperatures 20-32 °C and three for temperatures 38-45 °C. To estimate maximum ATP hydrolysis and Rubisco activation activities and the corresponding temperatures at which the maximum values are attained, second-to third-order polynomials and generalized additive models (GAM) were fitted to the experimental data using the gam function from the mgcv 1.8–24 package in R (Wood, 2017). The model that best fit the experimental data was selected based on the Akaike information criterion using the AIC function (Akaike, 1974) (Supporting Information Table S4). The selected model was fitted to each set of the biological replicates (Table 1) as well as the whole dataset (Supporting Information Table S3). T_max_ was calculated as the temperature corresponding to maximum activity. T_opt_ was calculated as the range where activity was maintained above 70% of maximum value. T_0.5_ was calculated as the temperature where activity dropped below 50% of maximum value.

**Table 1.** Temperature response of *in vitro* Rubisco activase (Rca) activity. Maximum rate of ATP hydrolysis and Rubisco activation and corresponding temperature (T_max_), optimum temperature range (T_opt_, above 70% activity) and temperature above the optimum at which 50% of the maximum activity remains (T_0.5_). Values were estimated from best-fit models that describe the *in vitro* temperature response of each Rca isoform (Supporting Information Table S4). The best-fit model was applied to each of three biological replicates with complete temperature response curves shown in Figure 1b. The first biological replicate was excluded from the fitting of the individual replicates because the temperature response of Rca activity was incomplete (determined only up to 32°C); this replicate is included in the model fitting to the complete temperature response data set (Supporting Information Table S3). Values show here are the mean ± standard error of the mean of values determined for individual replicates of temperature response curves (n=3).

| Rca Isoform | ATPase<sub>max</sub><br>(mol min<sup>-1</sup> mol Rca<sup>-1</sup>) | T<sub>max</sub><br>(°C) | T<sub>opt</sub><br>(°C) | T<sub>0.5</sub><br>(°C) |
| --- | --- | --- | --- | --- |
| 1β | 28.0 ± 5.3 | 36.7 ± 0.5 | 29.9 – 43.4 | 43.4 ± 1.1 |
| 8α | 35.9 ± 8.1 | 35.9 ± 0.7 | 28.4 – 43.1 | 45.6 ± 0.2 |
| 10α | 179.0 ± 36.8 | 42.6 ± 1.1 | 30.9 – 51.6 | 53.6 ± 0.7 |
| 10β | 183.4 ± 58.5 | 40.9 ± 1.1 | 28.5 – 52.1 | 53.5 ± 0.3 |
| Rca Isoform | Rubisco reactivation<sub>max</sub><br>(mol R<sub>A.S.</sub> min<sup>-1</sup> mol Rca<sup>-1</sup>) | T<sub>max</sub><br>(°C) | T<sub>opt</sub><br>(°C) | T<sub>0.5</sub><br>(°C) |
| 1β | 0.10 ± 0.03 | 29.5 ± 1.2 | 27.7 – 34.4 | 36.1 ± 1.5 |
| 8α | 0.09 ± 0.02 | 26.0 ± 2.9 | 22.1 – 36.0 | 41.0 ± 2.9 |
| 10α | 0.14 ± 0.01 | 34.5 ± 1.0 | 26.5 – 40.8 | 42.4 ± 0.2 |
| 10β | 0.19 ± 0.01 | 32.5 ± 0.4 | 24.0 – 40.9 | 43.3 ± 0.8 |

### Plant growth and sampling

*Vigna unguiculata* (L.) Walp. cv. IT97K-499-35 seeds (kindly provided by B.B. Singh) were germinated in 0.6 L Deepots (D40H, Stuewe & Sons) containing 1:1 (v:v) mixture of nutrient-rich compost (Petersfield Growing Mediums) and silver sand (horticultural grade, Royal Horticultural Society). Plants were grown in eight growth cabinets (MC1000, Snijders Labs) and watered as needed to soil saturation. Cabinets were maintained at day/night temperatures of 28/18 °C, 70% relative humidity and 400 ppm CO_2_ concentration (control conditions). A 12 h photoperiod of constant light of 850 µmol m^−2^ s^−1^ at canopy level was maintained via LED lighting (NS1 spectrum, Valoya). Four cabinets were randomly selected to apply a heat treatment starting at 13 days after sowing (das) including an intermediate day with day/night temperatures of 34/24 °C followed by 5 days of 38/28 °C (Supporting Information Fig. S3a). Data loggers (OM-EL_USB-2, Omega Engineering Ltd.) were placed at canopy level in each cabinet to record temperature and relative humidity every 15 minutes (Supporting Information Fig. S3b).

Twenty cowpea plants were grown in each cabinet and sampled or measured for different analyses following a randomised design (Supporting Information Fig. S4). All samples and measurements were taken from the central leaflet of the first trifoliate leaf. Leaf disc samples were taken 4-5 h into the light period, after which plants were cut above the unifoliate leaves to reduce shading of neighbouring plants. Separate plants were used for sample collection in subsequent time points. For non-destructive measurements, such as plant growth traits and chlorophyll content, the same individuals were measured at all time points.

Growth measurements were obtained at 15, 17 and 22 das (days 2 and 4 of the heatwave and on day 4 of the post-heatwave recovery period, Supporting Information Fig. S6). Leaf chlorophyll content was estimated using a chlorophyll concentration meter (MC-100, Apogee) by measuring 3 positions on the central leaflet for each plant. Plant height was measured using a ruler from the soil level to the apical meristem. Leaf length and width (widest part perpendicular to the length) were also measured with a ruler for the central leaflet only. Leaf thickness was measured with a digitronic calliper 110-DBL (Moore & Wright). Total above-ground biomass was determined on a separate set of plants by drying leaves and stems at 60 °C until constant weight.

Chlorophyll fluorescence (method below) measurements were conducted every day of the heatwave. Three 0.38 cm^2^ leaf discs were excised and quickly placed with the adaxial surface upwards on a single layer of wet filter paper on a thin plastic tray (<0.5 mm) and immediately transported to the fluorescence imager.

A thermal imaging camera (CAT S60 smartphone, Caterpillar) was used prior to sampling to assess T_leaf_. Each image was later analysed with FLIR Thermal Studio software (Teledyne FLIR LLC), providing the mean temperature across an area of the leaf surface (Supporting Information Fig. S5).

For RNA-seq analyses (method below), four 0.55 cm^2^ leaf discs were collected from 4 independent biological replicates per treatment per timepoint (days 1, 3 and 5 of heatwave) using a cork borer, then immediately snap frozen in liquid nitrogen and stored at −80 °C.

For protein and Rubisco analyses, four 0.55 cm^2^ leaf discs were excised as above and quickly placed with the adaxial surface upwards on a single layer of wet filter paper to maintain turgidity. The filter paper was contained in a thin plastic tray and positioned on the surface of the water in a circulating water bath in the artificial sunlight simulation rig (Taylor *et al*., 2022). PPFD at leaf-disc-level was 850 µmol m^-2^ s^-1^. Water bath temperature was maintained at 26 °C or 32 °C (±0.1 °C) for control or heat-treated plants, respectively, to mimic leaf temperatures in the growth cabinet. After 1 h incubation leaf discs were quickly blotted on absorbent paper to remove excess moisture, then immediately frozen in liquid nitrogen and stored at −80 °C.

For Rubisco reactivation in leaf extracts (LE), eight 0.55 cm^2^ leaf discs were sampled on day 5 of the heatwave. A different plant was used for each of the seven temperatures of the temperature response curve (20, 25, 32, 38, 40, 45 °C). Samples were quickly frozen in liquid nitrogen, then stored at −80 °C until further analysis.

### Chlorophyll fluorescence

Samples were analysed using a chlorophyll fluorescence imager (closed 800C FluorCam, Photon System Instruments). The imager contained a customized temperature-controlled sample surface area which comprised an aluminium plate with imbedded copper piping connected to a circulating temperature-controlled water bath (Optima TX150, Grant Instruments). The filter paper was in contact with sponges which acted as water reservoirs to ensure the paper remained wet during analysis (Supporting Information Fig. S7a). The temperature of the sample surface area and leaf discs were recorded using type k thermocouples and a data logger (RDXL4SD, Omega). Additional thermocouple wires were placed in contact with the sample surface area, and extra leaf discs, to record temperature throughout the analysis (Supporting Information Fig. S7a). Samples were first dark adapted for 1 h at 26 °C. After dark adaptation, the Fv/Fm protocol in the software (FluorCam 7, Photon System Instruments) was used to make measurements of the maximum quantum efficiency of photosystem II over a range of temperatures. The temperature of the leaf discs was used to assess when the measurement temperature had been reached and was stable. Once stable, the temperature was held for 3 min before the Fv/Fm measurement was taken.

The data for each set of three leaf discs from the same plant were averaged and plateauing Fv/Fm points at higher temperatures were removed before the remaining data were used to calculate T_crit_ (Supporting Information Fig. S7b) with the segmented function from the segmented package(Muggeo, 2017) in R. This function uses a previously fitted linear model to identify a breakpoint and fits two linear models before and after the breakpoint. The breakpoint in the Fv/Fm curve was taken as T_crit_ while the slopes of the two linear models were taken as m1 & m2 (Supporting Information Fig. S7b, c, d).

### Protein extraction, Rubisco activity and content

Proteins were extracted from leaf disc samples and Rubisco activity assays were performed in a randomised order, as described previously (Taylor *et al*., 2022; Ashton *et al*., 2024). The activation state of Rubisco was calculated as the ratio of initial to total activity. In short, aliquots of the leaf extract supernatant were added to an assay mix containing ^14^CO_2_ and RuBP to initiate assays that gave an estimate of Rubisco initial activity, whereas Rubisco total activity was measured after allowing a 3-minute incubation period of the leaf extract in the assay mix without RuBP to allow full carbamylation of Rubisco.

Aliquots of leaf extract supernatant were also used for determining Rubisco content via ^14^C-CABP (2-carboxy-D-arabinitol 1,5-bisphosphate) binding, as previously described (Ruuska *et al*., 1998; Ashton *et al*., 2024) and total soluble protein content, using the Bradford method (Bradford, 1976).

### Gene expressions analysis

Frozen leaf disc samples were ground to a fine powder in liquid nitrogen using a pestle and mortar. Total RNA was extracted from 20-30 mg tissue using a NucleoSpin™ RNA Plant Kit (Macherey-Nagel) including a DNase treatment, according to the manufacturer’s instructions. RNA concentration and quality were determined by spectrophotometry using a microplate reader and LVis plate (SPECTROstar Nano, BMG Labtech) (Supporting Information Table S5). Further QC analysis of RNA samples was performed by Novogene using a Qubit fluorometer (Invitrogen) and an Agilent 5400 fragment analyzer, followed by mRNA purification (using poly-T oligo-attached magnetic beads), library construction and 2 x 150 bp paired-end (PE150) sequencing using an Illumina NovaSeq 6000 (read depth of 30 million reads).

An initial filtering was applied to the raw data to remove reads containing adapters, containing N > 10% (N represents bases that cannot be determined) and low-quality bases (Phred < 5). Further read quality assessment and trimming was performed using the FastQC tool, version 0.12.1 (http://www.bioinformatics.babraham.ac.uk/projects/fastqc/) and cutadapt, version 4.2 (Martin, 2011). Low quality bases (Phred < 30) were trimmed from 3’ ends of each read, flanking N bases were removed and trimmed reads shorter than 50 bp were discarded (for sequencing and alignment statistics see Supporting Information Table S6).

Cowpea IT97K-499-35 reference transcriptome v1.2 (Lonardi *et al*., 2019; Liang *et al*., 2024) was downloaded from Phytozome 13 (Goodstein *et al*., 2012). Salmon, version 1.10.1 (Patro *et al*., 2017) was first used to index the reference transcriptome and then quantify expression of clean reads at the transcript level. To identify changes in the cowpea Rca transcripts TPM values were extracted from the Salmon Quant.sf output files for Rca, RbcS and HSP20 transcripts. Three transcripts are present for Rca1β (the first two result in the same protein, the third is missing some of the N domain) and TPM values for all three were combined.

Three transcripts exist for Rca8α (transcripts 2 and 3 are non-functional and have low expression) so here we only present data for the first transcript. For Rca10 transcripts 1, 3 and 4 are combined to give total expression for the alpha isoform and transcript 2 alone represents the beta isoform. The three transcripts for RbcS were also combined.

Global gene expression analysis was carried out in R version 4.4.1 (2024-06-14, Race for Your Life). Raw counts from Salmon were loaded into EdgeR 4.2.1 using tximport 1.32.0 and rtracklayer 1.64.0. Transcript data for each gene was combined at this stage so that all global analysis was performed at the gene-level. PCA plot was made using logCPM the PCAtools 2.16.0 to check that biological replicates group together and to look for outlying samples. Here we used the counts per million (CPM) to remove the variability due to library depth. The EdgeR trimmed mean of M-values (TMM) method was used for normalisation of read counts. The data was filtered using a CPM threshold to remove genes with zero or near zero expression. A generalised linear model was then fitted to the normalised and filtered data for each gene. Empirical Bayes quasi-likelihood F-tests were used to test whether genes were significantly differentially expressed between heat-treated and control plants on each day of the heatwave. To control for multiple testing, p-values were calculated using the false discovery rate (FDR) of Benjamini-Hochberg. Differentially expressed genes (DEGs) were then identified using the threshold values of a false discovery rate (FDR) of < 0.05 and a log2 (|fold change|) of >1.5.

Rca gene expression changes were validated via RT-qPCR (Supporting Information Fig. S11). For sampling and detailed methodology see Supporting Information Tables S7 & 8.

### Rca quantification via western blotting

To quantify the abundance of different Rca protein isoforms, an aliquot of supernatant resulting from Rubisco analysis was mixed with SDS-PAGE loading buffer and separated as previously described on Perdomo *et al*. (2018) on a 12% TGX Stain-Free gel (Bio-Rad). Gels were imaged to assess total protein before transfer to a nitrocellulose membrane using a dry blotting system (iBlot2, Thermo Fisher Scientific). Membranes were stained with REVERT Total Protein Stain for normalisation (Li-Cor Biosciences) and imaged prior to application of antibodies. To compare the abundance of α and β isoforms of Rca, an antibody with broad specificity for both isoforms was used (Feller *et al*., 1998,a gift from Michael Salvucci), with detection via a secondary fluor-tagged antibody (IRDye800CW, Li-Cor Biosciences, RRID:AB_1660973). A dilution series of a pooled sample was run on every gel alongside the samples to verify antibody detection was within the linear range. To quantify Rca8α, a more specific antibody was used (Abcalis) in place of the broad specificity antibody. The anti-Rca8α specific antibody was generated using phage display with the peptide selected targeted against KRGAFYGKAAQQINVP (amino acid residue 376-391), as described in Bloemers & Carmo-Silva (2024). All blot images were obtained using an Odyssey FC (Li-Cor Biosciences). Empiria studio software (version 2.3.0.154, Li-Cor Biosciences) was used for image data analysis and normalisation with the total protein stain, with values determined as signal intensity.

### Characterization of Rca thermal optima in leaf extracts

Rubisco reactivation by Rca in leaf extracts was determined as described in Carmo-Silva & Salvucci (2011). Eight leaf discs per plant were sampled from the youngest fully expanded leaf on day 5 of the heatwave. Separate samples were used for assaying each of the seven temperatures of the temperature response curve shown in Figure 4, and eight plant sample replicates were used per temperature. Samples were stored at −80 °C until further analysis. One replicate of samples from one cabinet at control and one cabinet at high temperature was used in each day of Rca assays, with samples allocated randomly to each assay temperature. Supporting Information Fig. S14 illustrates the method which uses leaf extracts as the source of both Rubisco and Rca, incubates sub-aliquots with different assays mixes, and then carries out a two-stage assay very similar to the assay with the purified proteins. This enables determination of the capacity of a population of Rca, drawn from plants grown in different conditions, to reactivate Rubisco. Model fitting and temperature parameter (T_max_, T_opt_ and T_0.5_) calculations were conducted as described for the *in vitro* Rca activity assays above. From the same samples, aliquots of leaf homogenate were used for leaf chlorophyll content determination as described previously in Sales *et al*. (2022) and Taylor *et al*. (2020), according to equations by Lichtenthaler (1987). Aliquots of the desalted extract were used for total soluble protein (TSP) content determination using the Bradford method (Bradford, 1976). Results are presented in Supporting Information Table S3.

**Table 2.**
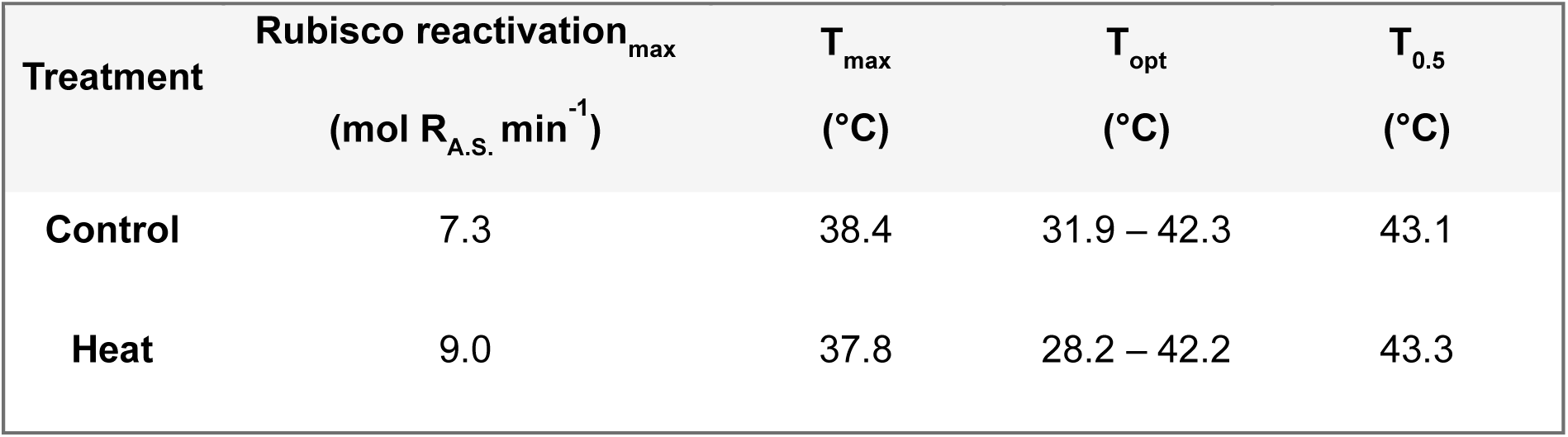
Temperature response of Rubisco activase (Rca) activity in leaf extracts. Maximum rate of Rubisco activation and corresponding temperature (T_max_), optimum temperature range (T_opt_, above 70% activity) and temperature above the optimum at which 50% of the maximum activity remains (T_0.5_). Values were estimated from best-fit models that describe the temperature response of the pool of Rca isoforms extracted from leaves of control and heat-treated cowpea on day 5 of the heatwave (Supporting Information Table S9). The best-fit model for each treatment was applied to combined replicates shown in Figure 4 (n=6-8).

### Data analyses

Geneious 2024.0.7 was used to generate protein alignments depicted in Figure 1a. Data was processed using R 4.3.1 and RStudio 2024.09.0. Graphs were prepared using the ggplot 2 (Wickham, 2017), EnhancedVolcano (Blighe *et al*., 2018), VennDiagramm (Chen, 2022), ggpubr (Kassambara, 2023) and patchwork (Lin Pedersen, 2024) R packages. Outliers were detected with the Outliers package using Tukey’s fences method, where outliers are defined as extreme values that are 1.5 times the interquartile range (1.5 IQR) below the first quartile or 1.5 IQR above the third quartile. Box plots show medians and the first and third quartiles (25^th^ and 75^th^ percentiles), and whiskers extend from the hinge to the largest or smallest value. Symbols represent individual data points (technical or biological replicates defined for each plot either on methods or figure legend). Bar plots show means and error bars represent standard error. For statistical analysis, normality of data was assessed using the Shapiro-Wilk test and equality of variances using the Levene test. For measurements on multiple days, two-way ANOVA tests were applied to determine if there were significant day, treatment or interactive effects, followed by post-hoc Tukey tests for multiple comparisons with Bonferroni correction. Compact letters display (cld) were generated after the post-hoc pairwise comparisons using packages mutlicomp (Bretz *et al*., 2016) and multicompView (Graves *et al*., 2024). In cases where there was a slight violation of normality or homogeneity of variance when data between different days was compared, two-way ANOVA was still chosen as the most robust method. If normality and variance were severely violated, data was log transformed, and two-way ANOVA was applied to the transformed data provided assumptions were not severely violated in the transformed data. For measurements taken at a single timepoint (growth parameters), t-test was used to determine statistical significance between treatments. Statistical analysis results and full datasets are provided in the Source Data file.

**Figure 1.**
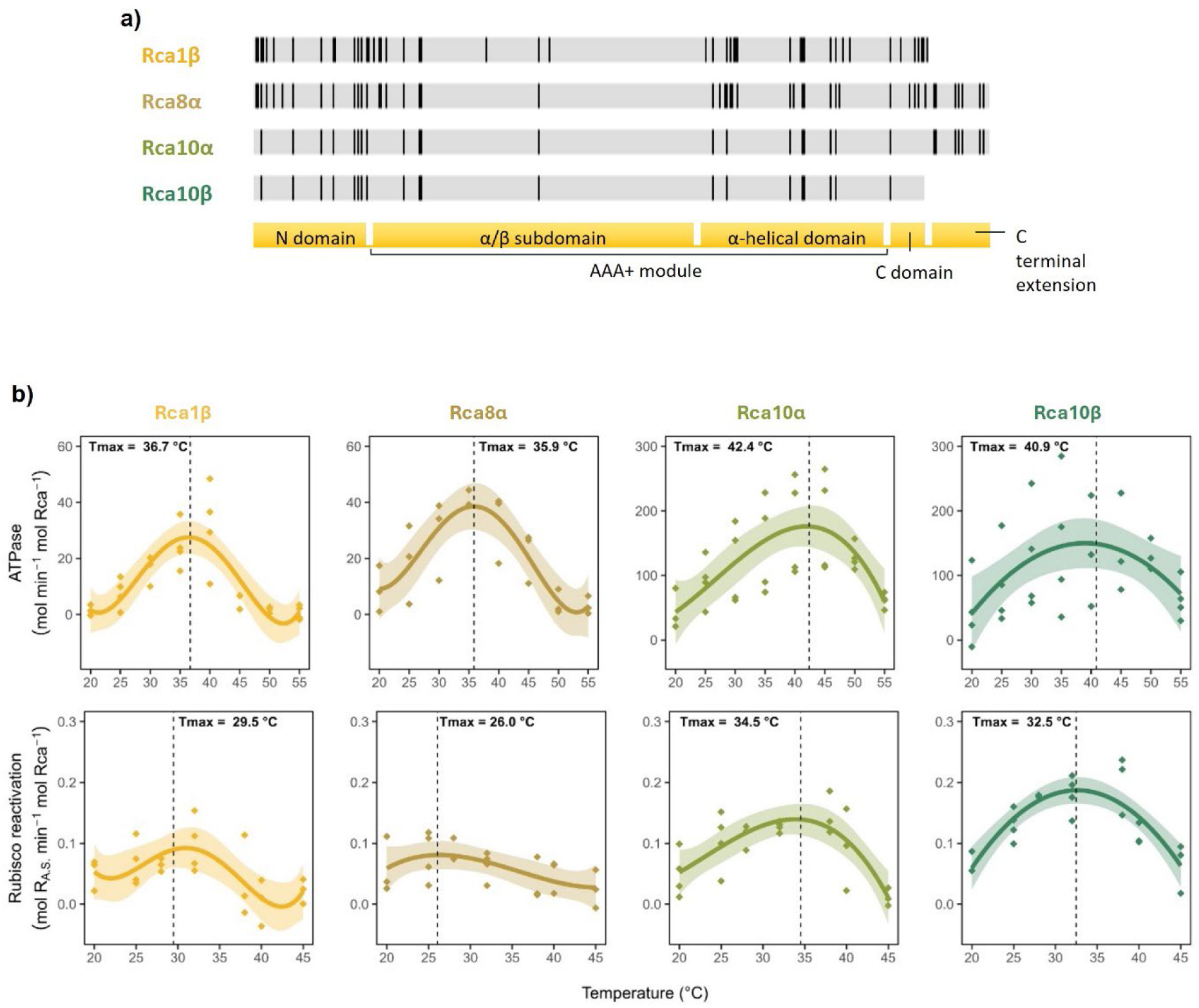
Temperature response of cowpea Rca isoforms in vitro. a) Amino acid residue alignment of the four cowpea Rca isoforms encoded by genes located on chromosomes 1, 8 and 10. Black vertical lines indicate differences in sequence among the genes. The following corresponding regions of the protein are shown: N-domain, α/β subdomain, α-helical domain, C-domain, and C-terminal extension. b) Temperature response of ATP hydrolysis and Rubisco activation by cowpea Rubisco activase isoforms *in vitro*. Rate of ATP hydrolysis and Rubisco reactivation by recombinantly produced purified cowpea Rca isoforms Rca1β, Rca8α, Rca10α and Rca10β. Assays were performed at the indicated temperatures using 1 μM purified Rca and 5 mM ATP. Activation of pre-inhibited Rubisco used 5 μM Rubisco active sites (R_A.S._) in the ER form (1:5 Rca:R_A.S_). Rubisco activation is presented relative to maximum Rubisco activity, determined as the Rubisco activity when Rca is allowed to reactivate inhibited Rubisco for 5 min. Values represent biological replicates (independent purifications; n=3-4 biological replicates). Lines represent the best fit for each enzyme (selected by the Akaike information criterion) and shaded areas denote the 95% confidence interval for each fit (Supporting Information Table S4). Full sample data are provided in the source data file.

## Results

### Cowpea contains four Rubisco activase isoforms

The cowpea genome contains three genes that encode Rca. The gene on chromosome 1 (*Vigun01g219300)* encodes an Rca β, the gene on chromosome 8 (*Vigun08g150700)* an Rca α, and the gene on chromosome 10 (*Vigun10g051600)* encodes both Rca α and β isoforms via alternative splicing (Figure 1a). The gene on chromosome 8 also encodes a truncated version of Rca, however this would not produce a canonical, fully functional Rca as it has a premature stop codon before the α-helical domain which is necessary for the recognition of Rubisco by Rca (Portis *et al*., 2008). Rca isoforms in cowpea exhibit minor sequence differences, primarily in the N-terminal region (the flexible domain critical for Rubisco binding) (Stotz *et al*., 2011; Shivhare *et al*., 2019, p. 20) and the α-helical domain (Supporting Information Fig. S1). These variations may contribute to functional diversity of Rca in cowpea, potentially allowing for fine-tuning of responses to different environmental conditions.

### Cowpea Rca isoforms differ in thermal optima of activity *in vitro*

Previous studies have shown the potential for intraspecies variation of Rca thermal optima (Degen *et al*., 2020). With four distinct isoforms in cowpea and its known ability to grow in warm (20-37 °C) temperatures (Barros *et al*., 2021), recombinant versions of each isoform were produced for *in vitro* characterisation of cowpea Rca thermal optima (Figure 1b). Both enzyme activities were assessed, i.e., the ability to reactivate inhibited cowpea Rubisco and the ATPase activity that drives reactivation (Robinson & Portis, 1989). Rubisco reactivation by Rca was calculated after accounting for the spontaneous reactivation of Rubisco, expressed as a percentage of maximum Rubisco activity. The spontaneous reactivation of Rubisco, although insignificant at lower temperatures, increases with temperature (Supporting Information Fig. S2) as the Rubisco catalytic site becomes more flexible. For all the Rca isoforms, the temperature corresponding to the peak of activity (T_max_), the range of temperatures at which activity was maintained above 70% of the maximum value (T_opt_) and the temperature at which activity drops below 50% of the maximum (T_0.5_) were consistently higher for ATPase activity than for Rubisco reactivation activity (Table 1). A higher sensitivity of Rubisco reactivation compared to ATP hydrolysis has been previously reported and may be due to the interaction with Rubisco being the key step impacted by elevated temperatures in Rca (Degen *et al*., 2020). It has also been shown that while ATP hydrolysis can be catalysed by Rca oligomers consisting of three subunits, Rubisco reactivation requires oligomers containing more than three subunits (Keown *et al*., 2013; Hazra *et al*., 2015).

The two Rca isoforms encoded by *Rca10* were faster at hydrolysing ATP and reactivating Rubisco, had higher thermal maxima (T_max_), broader thermal optima (T_opt_), and showed thermal sensitivity (T_0.5_) at warmer temperatures compared to the other two isoforms (Figure 1b;Table 1). T_max_, T_opt_, and T_0.5_ calculated for both ATP hydrolysis and Rubisco reactivation were generally similar for Rca1β and Rca 8α (Table 1), but Rca1β exhibited the lowest T_0.5_ for reactivation activity (36.1 ± 1.5 °C). Up to 34 °C, all Rca isoforms maintained at least 70% of maximum Rubisco reactivation activity. Rca10α exhibited the highest temperature corresponding to the peak of activity for both ATP hydrolysis (T_max_ = 42.6 ± 1.1 °C) and Rubisco reactivation (T_max_ = 34.5 ± 1.0 °C). T_max_ values for Rca10α and Rca10β were at least 3 °C higher for both ATP hydrolysis and Rubisco reactivation relative to Rca1β and Rca8α (Table 1). Combined, the results show that the cowpea Rca10α and Rca10β isoforms have faster rates, higher thermal maxima, broader thermal optima, and are less sensitive to warmer temperatures than Rca1β and Rca8α.

### A 5-day heatwave of +10 °C causes mild heat stress in cowpea

To investigate heat-induced changes in cowpea Rca *in planta*, young plants were exposed to a 5-day heatwave consisting of an increase of 10 °C during both day and night. Plants were cultivated for two weeks at 28/18 °C day/night, then in half of the controlled environment cabinets the temperature was increased to 34/24 °C day/night for one day and subsequently to 38/28 °C day/night for five days (Figure 2a, Supporting Information Fig. S3, 4). The intermediate day aimed to resemble a gradual increase in temperature in field conditions. Plants were well watered throughout the heatwave as the focus was on the impact of heat on Rca and we wanted to avoid confounding effects of water deficit on CO_2_ diffusion and assimilation. Cabinet and leaf temperatures were monitored throughout the experiment to ensure the plants were under heat stress conditions, with heat-treated plants showing a 6 °C increase in leaf temperature to 32 °C (Figure 2a, Supporting Information Fig. S5). Thus, plants experiencing the heatwave had leaf temperatures within the T_opt_ of Rca activity.

**Figure 2.**
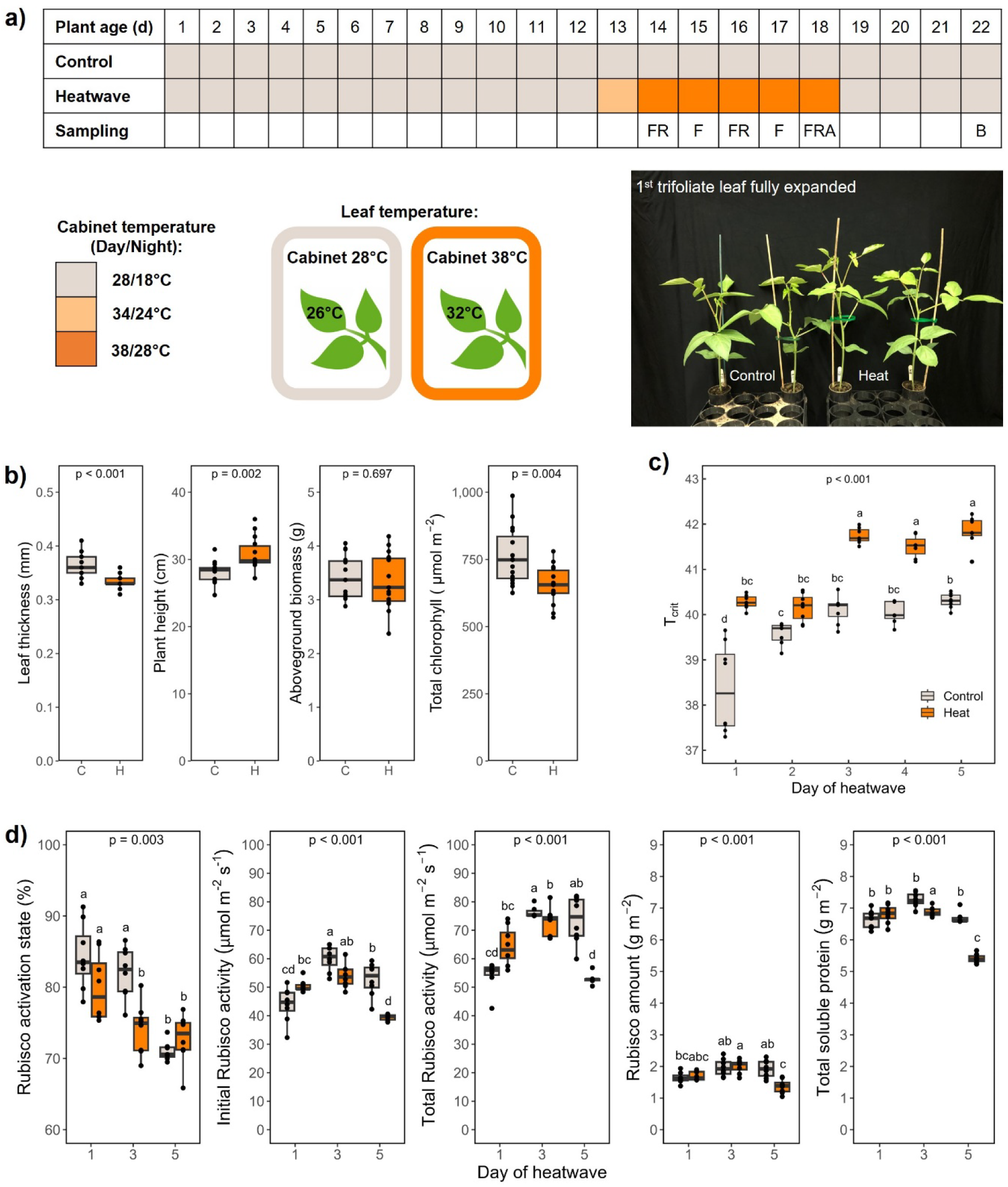
Cowpea response to a heatwave in controlled environment. a) Experimental design, heatwave and sampling times. Plants were grown at 28/18 °C day/night for 13 days, then exposed to an intermediate heat treatment for a day at 34/24 °C day/night before being exposed to a 38/28 °C day/night heatwave for five days. Sampling: (F) photosystem II maximum efficiency (Fv/Fm) was measured daily during the heatwave; (R) leaf discs for RNA-seq analysis, total soluble protein (TSP), Rubisco activity and amount were sampled on days 1, 3 and 5 of the heatwave; (A) samples for Rca temperature response in leaf extracts were taken on day 5 of heatwave; (B) growth parameters and biomass were measured on day 22 of plant growth, four days after the end of the heatwave. Cabinet and leaf temperature were monitored daily. b) Leaf and whole plant growth parameters were measured four days after the end of the heatwave (n=15). Relative chlorophyll content was determined using a concentration meter based on leaf reflectance. c) T_crit_ was calculated for each day of the heatwave as the breaking point where the two Fv/Fm slopes meet (Supporting Information Fig. S7). d) Rubisco activation state (n=7-8), (b) initial (n=7-8) and (c) total activity per leaf area (n=6-8), Rubisco amount (n=7-8) and total soluble protein (TSP; n=6-8) were measured in leaf extracts on days 1, 3 and 5 of the heatwave. P-values were calculated using two-way ANOVA followed by Tukey post-hoc tests. Full sample data are provided in the source data file.

Plant growth traits were determined four days after the heatwave, after plants were allowed to recover, and showed that the 5-day heatwave of +10 °C had only a mild impact on cowpea (Figure 2b). Exposure to elevated temperature resulted in decreased leaf thickness and increased plant height; however, no effect on aboveground dry biomass was observed. Total chlorophyll also decreased in leaves that had been exposed to heat when determined four days after the heatwave (Figure 2b). Measurement of leaf and plant traits during the heatwave suggested an initial increase in the rate of cowpea leaf and plant growth (day 2, Supporting Information Fig. S6), but only minimal differences between control and heat at the end of and after the heatwave.

As a proxy for photosynthetic activity, photosystem II (PSII) maximum efficiency (Fv/Fm) was monitored throughout the heatwave (Supporting Information Fig. S7). Changes in Fv/Fm indicate modifications in the functionality of PSII reaction centres, with a decrease indicating photooxidative damage and reduced photochemistry quantum yield (Murchie & Lawson, 2013). Fv/Fm data collected from plants throughout the heatwave enabled determination of T_crit_ ( Figure 2c), which corresponds to the temperature at which chlorophyll *a* fluorescence rises rapidly in response to temperature due to thermal damage in PSII, and is used as an indicator of photosynthetic thermal stability (Perez & Feeley, 2020; Posch *et al*., 2022). In heat-treated cowpea, Fv/Fm did not decline compared to control plants (Supporting Information Fig. S7e), indicating that no irreversible damage was caused to PSII reaction centres due to heat stress. In fact, heat-treated plants showed a progressive increase in the temperature at which Fv/Fm decreased, resulting in a significantly higher T_crit_ after 3 days, denoting increased PSII thermal stability (p < 0.001, Figure 2c). These results indicate a dynamic response that allows for PSII efficiency to adapt to warmer conditions.

### Heat stress induces a decline in Rubisco activation state

In contrast to T_crit_, the activation state of Rubisco decreased with heat treatment (Figure 2d). Importantly, a decline in Rubisco activation was observed from the beginning to the end of the experiment as leaves aged. This decline was observed earlier in heat-treated plants (day 3) compared to control plants (day 5), indicating that heat stress accelerated this decline in Rubisco function. Rubisco activation state, defined as the ratio of initial to total Rubisco activity, reflects the enzyme’s operational efficiency under different conditions. Initial activity represents the enzyme’s immediate functionality upon extraction, while total activity reflects its potential after full carbamylation (Ashton *et al*., 2024). Although Rubisco activation state was similar on day 5, the initial and total activity was significantly lower in heat-treated compared to control plants (p < 0.001, Figure 2d). This decrease in Rubisco activity on day 5 of the heatwave was associated with decreased Rubisco amount and TSP (Figure 2d), as no significant differences were detected in Rubisco specific activity, expressed per quantity of Rubisco protein (Supporting Information Fig. S8).

### Heat stress induces the expression of thermotolerant Rca isoforms

Potential heat-induced changes in gene expression of *Rca* isoforms were assessed through RNA sequencing analysis (RNA-seq) on days 1, 3 and 5 of the heatwave. A principal component analysis (PCA) distinguished gene expression changes caused by heat stress and those resulting from leaf aging (Figure 3a, Supporting Information Fig. S9). Heat-induced differential gene expression occurred from the first day of the heatwave, with both up and down-regulation of genes evident in heat-treated plants compared to control conditions throughout the duration of the heatwave (Supporting Information Fig. S9). Heat stress was confirmed by upregulation of the heat shock protein *HSP20* (Song *et al*., 2017) from day 1 of the heatwave (Figure 3b). The heatwave also decreased expression of the Rubisco small subunit (*RbcS*), which is likely associated with the lower Rubisco amounts (Figure 3c).

**Figure 3.**
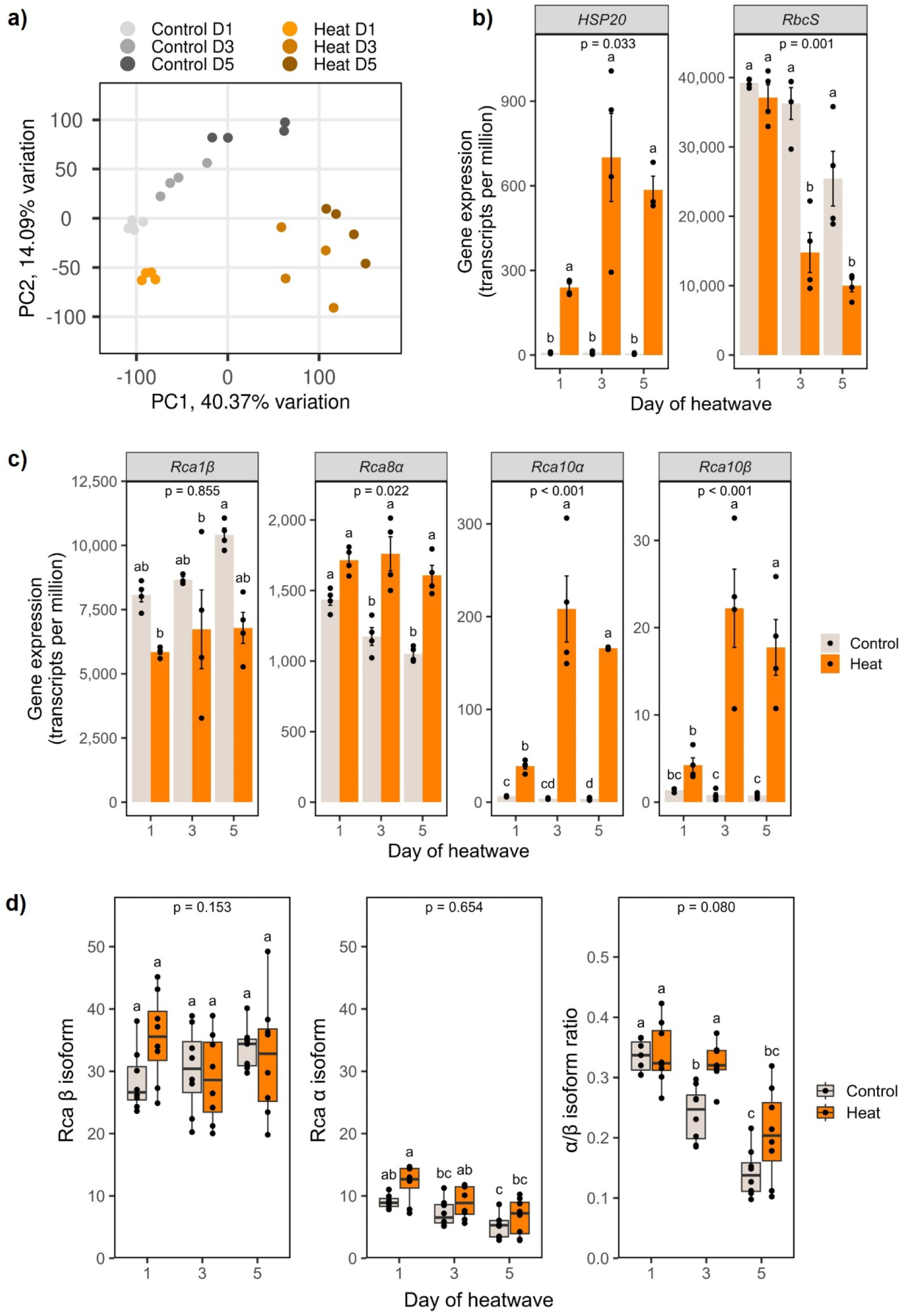
Differential gene expression and Rca protein abundance in cowpea during a heatwave. a) Principal components (PCA) plot of gene expression in leaves of heat-treated and control cowpea plants on days 1, 3 and 5 of the heatwave. b) Expression levels of the heat shock protein 20 (HSP20) and the small Rubisco subunits (RbcS) were used as reference. Bar plots represent means and error bars standard error. c) Expression levels of the four cowpea Rca isoforms in control and heat-treated plant on days 1, 3 and 5 of the heatwave (n=4). d) Protein abundance of Rca isoforms in leaves of control and heat-treated plants on days 1, 3 and 5 of the heatwave (relative units). Rca α and β isoforms were detected using an anti-Rca antibody and normalised to total protein. The size difference between the α (45 kDa) and β (41 kDa) isoforms enabled the identification as separate bands, and the estimation of the α/β ratio. P-values were calculated using two-way ANOVA followed by Tukey post-hoc tests. Samples were run in separate gels and normalised to total protein via reversible staining of the membrane. Blot images and full sample data are provided in the source data file.

Under control temperature conditions, *Rca1β* was found to be the most abundant Rca transcript, followed by *Rca8α*, while *Rca10α* and *Rca10β* were expressed at much lower levels (Figure 3c). Of the hundreds of up and downregulated genes (Supporting Information Fig. S9), volcano plots showed that *HSP20* and *Rca10* were both upregulated on days 1, 3, and 5 of the heatwave (Supporting Information Fig. S10). Heat-treated plants showed a (non-significant) tendency for lower *Rca1β* expression, whereas *Rca8α*, *Rca10α* and *Rca10β* were all upregulated during the heatwave (Figure 3c). It is noteworthy that despite the upregulation of both *Rca10* transcripts, their expression levels remained much lower than *Rca1β* and *Rca8α* (Figure 3c). Changes in *HSP20* and the four *Rca* transcript levels were also confirmed via RT-qPCR (Supporting Information Fig. S11). Moreover, promoter analysis revealed multiple stress responsive elements in *Rca1β and Rca8α*, while *Rca10* contains two heat stress response elements and has fewer other stress responsive elements (Supporting Information Fig. S12).

To investigate if changes in gene expression are reflected at protein level, the abundance of leaf Rca protein was quantified via immunoblotting (Figure 3d). A pan-Rca antibody that reacts with all isoforms of Rca and enables quantification of Rca α and Rca β isoforms, as these run separately in gel electrophoresis due to the difference in molecular weight, was used to identify changes in abundance between α and β isoforms (Perdomo *et al*., 2018). Whilst differential gene expression was observed, changes in Rca protein were less pronounced. As *Rca10β* transcript levels are almost negligible compared to *Rca1β*, it is reasonable to infer that the β isoform protein is predominantly attributable to *Rca1β* expression. Expression of *Rca8α* is also likely to produce most of the α isoform protein. Rca β remained unchanged during the heatwave, while the α isoform decreased from day 1 to day 5 but was not significantly different between control and heat-treated plants (Figure 3d). Protein abundance of Rca8α was also determined using a specific antibody that only reacts with this isoform (Bloemers & Carmo-Silva, 2024). Rca8α showed a near identical pattern to the pan-Rca antibody for Rca α with a non-significant tendency for increased abundance in heat treated plants compared to the control (Supporting Information Fig. S13a). A significant increase in the α/β ratio was however detected in day 3 of the heatwave, which may indicate differences in the relative abundance of Rca isoforms (Figure 3d). Due to their very low abundance, Rca10α & Rca10β protein isoforms were undetected in leaf extracts with Rca10 specific antibodies (Supporting Information Fig. S13b).

### Heat-treated plants maintain Rca rate and thermal optimum

To test the hypothesis that heat-induced changes in the relative abundance of Rca isoforms are reflected in activity, we characterised the temperature response of Rubisco reactivation by Rca in protein extracts from leaves sampled on day 5 of the heatwave (Figure 4). Leaf extracts were desalted to produce the free Rubisco apoenzyme (uncarbamylated form), which was subsequently inhibited through incubation with RuBP (Carmo-Silva & Salvucci, 2011). The inhibited Rubisco in the leaf extract was then used to determine the rate of reactivation by the Rca holoenzyme via removal of RuBP from uncarbamylated Rubisco catalytic sites, at a range of temperatures (Supporting Information Fig. S14). Concurrent measurement of spontaneous reactivation of Rubisco in the absence of ATP, which is required for Rca activity, enabled calculation of the spontaneous (Supporting Information Fig. S15) and Rca-mediated rate of Rubisco reactivation (Figure 4).

**Figure 4.**
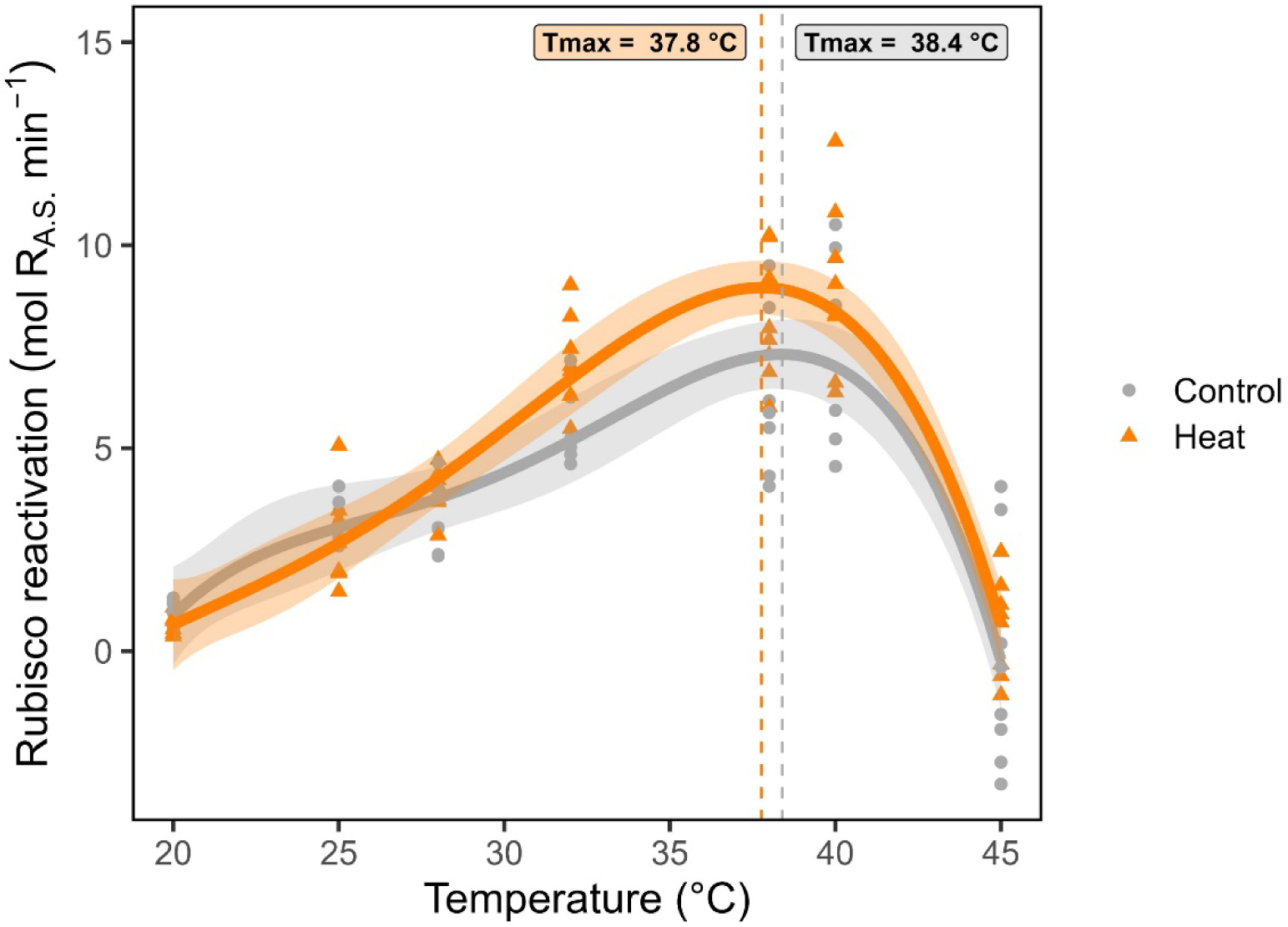
Temperature response of Rca activity in leaf extracts of control and heat-treated cowpea plants. Leaf discs were sampled on day 5 of the heatwave. The Rubisco and Rca contained in the leaves of each plant were extracted, Rubisco was inhibited by binding of RuBP to decarbamylated catalytic sites in the absence of CO_2_ and Mg^2+^, then reactivation by Rca was measured at varying temperatures by comparing an assay in presence of ATP and ATP-regenerating system with the negative control assay in the absence of ATP. The rate of Rubisco reactivation was calculated from measurements of Rubisco activity at 30 and 90 s after the start of the assay and expressed relative to maximum Rubisco activity in the same leaf extracts (as described in Supporting Information Fig. S14). Symbols represent biological replicates (leaf extracts from independent plant samples; n=8). Lines represent the best fit for each group of plants (selected by the Akaike information criterion) and coloured areas denote the 95% confidence interval for each fit (Supporting Information Table S9). Full sample data are provided in the source data file. Rubisco reactivation by Rca tended to be faster in heat-treated compared to control plants, however this increase was not statistically significant (*p > 0.05*, Figure 4). Heat-treated plants exhibited similar thermal optima of Rubisco reactivation by Rca to control plants at approximately 38 °C (Figure 4). T_opt_ and T_0.5_ were also similar between control and heat-treated plants (Table 2), indicating that the large fold-change in *Rca10* transcript abundance (Figure 3c) and the small change in heat-induced protein Rca isoform ratio (Figure 3d) did not alter the temperature profile of Rubisco reactivation by Rca in the leaves of heat-treated plants (Figure 4). Interestingly, the T_max_, T_opt_ and, to a lesser extent, T_0.5_ of Rubisco reactivation by Rca were higher in leaf extracts (Table 2) than the respective values measured for each individual Rca isoform purified after recombinant expression (Table 1).

## Discussion

### Cowpea Rca isoforms differ in thermal optima

We investigated the temperature response of Rubisco activase in cowpea, a vital crop for sub-Saharan Africa that is adapted to warm temperatures, yet susceptible to the increasing pressures of global warming. Several studies demonstrated that Rca thermolability limits photosynthesis at elevated temperatures (Feller *et al*., 1998; Crafts-Brandner & Law, 2000; Salvucci *et al*., 2006; Perdomo *et al*., 2017), making it a target for enhancing crop yield under elevated temperatures. In this study, we identified and characterised four Rca isoforms in cowpea: Rca1β, Rca8α, Rca10α and Rca10β. *In vitro* analysis across a range of temperatures revealed a higher thermal optimum for Rca10α and Rca10β for both ATPase and Rubisco reactivation activities (Figure 1b). The most abundant Rca1β and Rca8α isoforms in cowpea leaves (Figure 3) both exhibited lower thermal optima and thermal maxima, as well as slower rates of catalysis for both activities compared to the isoforms encoded by *Rca10* (Table 1).

Differences in thermal optima among isoforms have been reported for other species. In rice, the α isoform is more thermotolerant while in wheat one of the two β isoforms exhibits a higher thermal optima (Scafaro *et al*., 2018; Shivhare *et al*., 2019; Degen *et al*., 2020). For spinach, the β isoform is more sensitive to heat stress, with Rca β homo-hexamers displaying a lower thermal midpoint compared to Rca α homo-hexamers or hetero-hexamers comprising both α and β isoforms (Keown & Pearce, 2014). Notably, for all Rca isoforms, the T_max_ for ATPase activity is higher than for Rubisco reactivation, potentially due to differential oligomeric requirements for each activity. The proposed mechanism for Rca-mediated Rubisco reactivation involves threading Rubisco through the AAA+ pore, akin to the method employed by AAA+ ATPases that adjust the conformation of misfolded proteins (Bhat *et al*., 2017; Houry *et al*., 2017; Waheeda *et al*., 2023). This mechanism requires hexameric Rca for Rubisco reactivation, whilst ATP hydrolysis can occur with three Rca subunits forming an oligomer (Keown *et al*., 2013; Hazra *et al*., 2015).

The maximum rates of both ATP hydrolysis and Rubisco reactivation were higher for the Rca10 isoforms compared to Rca1β and Rca8α (Table 1). Similar differences among Rca isoforms were observed in rice and wheat but only for ATP hydrolysis, with the more thermostable Rca isoforms exhibiting lower Rubisco reactivation maxima compared to the less thermostable isoforms (Scafaro *et al*., 2016, 2019a; Shivhare & Mueller-Cajar, 2017; Degen *et al*., 2020). In cowpea, we show that the increased thermostability of Rca10 isoforms occurs without a penalty in Rubisco reactivation rate. The broader T_opt_ of Rca10α and Rca10β spans and extends above the T_opt_ of Rca1β and Rca8α (Table 1). Thus, Rca10α and Rca10β appear to be ideal candidates to maintain CO_2_ assimilation by Rubisco both at optimal and supra-optimal temperatures. Curiously, the promoter regions of both Rca1β and Rca8α contain multiple stress responsive elements while Rca10α and Rca10β have only two heat response elements (Supporting Information Fig. S12), suggesting an engineering approach may be required to alter the Rca isoforms predominantly involved in maintaining Rubisco activity *in planta*.

### Cowpea responds dynamically to a 5-day 38/28 °C heatwave

A 5-day heatwave at the vegetative growth stage (10 °C increase to 38/28 °C day/night) is considered moderate heat stress for most cowpea cultivars (Lonardi *et al*., 2019; Barros *et al*., 2021). Plants were well-watered to avoid confounding effects of water deficit limiting the diffusion of CO_2_ to Rubisco sites (Carmo-Silva *et al*., 2012). The leaf temperature only increased to 32 °C (Figure 2a), plant growth parameters were only mildly impacted, and the aboveground dry biomass remained unaffected (Figure 2bc). Notably, it has been shown that a heatwave during the vegetative stage can have a delayed impact on cowpea yield (Mohammed *et al*., 2024), which would not necessarily be expressed as reduced biomass at the end of the heatwave. T_crit_, an indicator of thermal lability of PSII, was higher on day 5 of the 38/28 °C heatwave (Figure 2c) suggesting some capacity for PSII thermal acclimation (Perez & Feeley, 2020). In wheat, increased T_crit_ after exposure to elevated temperature in field and controlled environment experiments correlated with a dynamic response of PSII to heat stress (Posch *et al*., 2022). On day 5 of the 38/28 °C heatwave, cowpea plants had decreased Rubisco activities and amount, lower leaf thickness, total chlorophyll and TSP (Figure 2b), suggesting accelerated leaf senescence. Similar results were observed in heat-treated wheat (Qu *et al*., 2023), which could have a delayed impact on biomass production.

### Thermotolerant Rca isoforms increase but remain low under heat

At optimal temperatures, *Rca1β* was the isoform most abundantly expressed, with *Rca8α* following, while *Rca10α* and *Rca10β* were only marginally expressed (Figure 3c). Under heat stress, *Rca8α*, *Rca10α* and *Rca10β* expression was upregulated, however *Rca10* transcript levels remained very low compared to *Rca1β* and *Rca8α*. The corresponding protein abundance was only marginally higher for Rca8α and remained undetectable for Rca10α and Rca10β (Supporting Information Fig. S13; Figure 3d). The Rca α/β ratio decreased as the leaves aged, and on day 3 of the heatwave heat-treated plants maintained higher α/β ratio than control pants (Figure 3d). Variation in Rca α/β ratio under optimal and supra-optimal temperatures may reflect differences in thermostability and regulatory mechanisms (Zhang & Portis, 1999; Wang *et al*., 2010; Degen *et al*., 2021). While Rca α is subject to redox regulation via thioredoxin-f (Zhang & Portis, 1999; Shivhare *et al*., 2019; Kim *et al*., 2020) and tends to be more sensitive to inhibition by ADP (Zhang & Portis, 1999; Scafaro *et al*., 2019b; Amaral *et al*., 2024a), variation also exists in the properties of Rca β isoforms (Scafaro *et al*., 2019a; Perdomo *et al*., 2019).

Changes in relative abundance of Rca isoforms under heat stress are species-specific (Yamori & von Caemmerer, 2009). In *Arabidopsis*, the two isoforms are comparable in thermotolerance and both increase with heat maintaining a 1:1 ratio (Ristic *et al*., 2009). Conversely, in rice and wheat, Rca isoforms differ in temperature response and heat induces upregulation of the thermotolerant variants (Scafaro *et al*., 2018, 2019a; Degen *et al*., 2021). While the same was observed for cowpea in the present study, the protein abundance of the thermostable Rca isoforms remained below detection levels in plants exposed to the 5-day 38/28 °C heatwave. Overexpression of thermostable Rca via genetic engineering led to enhanced photosynthesis and yield (Kurek *et al*., 2007; Shivhare & Mueller-Cajar, 2017; Scafaro *et al*., 2018; Degen *et al*., 2020). Increasing Rca thermostability, and to a lesser extent upregulation of its quantity, have been shown to ameliorate the reduction in Rubisco activation state caused by elevated temperatures (Kurek *et al*., 2007; Scafaro *et al*., 2018, 2019a; Kim *et al*., 2020). The results presented here suggest that increasing Rca10α and Rca10β abundance in cowpea, while possible in longer or hotter heat conditions, would likely require an engineering approach.

### Rubisco reactivation by Rca in cowpea remains unaltered under heat

The temperature response of Rubisco reactivation in leaf extracts of plants exposed to the 5-day 38/28 °C heatwave showed a similar pattern and comparable thermal optima to control plants (Figure 4). Scafaro *et al*. (2019a) reported increases Rubisco reactivation rate and T_0.5_ of Rca in leaf extracts from heat-treated wheat. In cowpea, the minor increase in rate was statistically insignificant and T_opt_ and T_0.5_ were comparable between control and heat-treated plants, with a slight increase of the T_opt_ range in heat-treated plants (Table 2). These results indicate that the heatwave in the absence of water deficit caused only mild heat stress, allowing plants to respond by adjusting isoform ratios in the leaf while preserving the Rca rate and temperature response.

Interestingly, the T_opt_ upper bound determined in leaf extracts was 42 °C for control and heat-treated plants, much higher than the T_leaf_ in heat-treated plants (Table 2, Figure 2a). The maximum Rca rate in leaf extracts (Table 2) was also higher than the Rubisco reactivation rate of each isoform measured *in vitro* (Table 1). Similar results were observed in rice and wheat but only for the more thermostable isoforms (Scafaro *et al*., 2016, 2019a). Higher Rca rate in leaf extracts may be attributed to several possible factors: differences in structural conformation of recombinant Rca, post-translational modifications that regulate Rca activity in the leaf and are absent in recombinant proteins, hetero-hexamers comprising different isoforms *in planta* exhibiting higher thermal optima, or interaction of Rca with other components in the chloroplast. Chen *et al*. (2010) reported dynamic and reversible binding of rice Rca to the thylakoid membrane while it is active in remodelling the conformation of inhibitor-bound Rubisco. Binding of Rca to the thylakoid membrane has also been reported during heat stress although its binding to the chloroplast ribosomes was proposed to have a protective role (Rokka *et al*., 2001). Moreover, heat stress has been shown to induce association of Rca with β-subunit of chaperonin-60 (cpn60β), a potential heat shock molecular chaperone (Salvucci, 2008). Thus, while *in vitro* analyses are useful to compare the properties of the individual isoforms, *in vivo* analyses as shown here for cowpea enable assessment of the overall Rca response within the chloroplast stroma.

## Conclusion

Cowpea is a vital source of protein for sub-Saharan Africa, and depite being warm-adapted, it is threatened by future heatwaves, which are likely to increase in intensity, duration, and frequency. Here, we identified two isoforms of Rca that have higher thermal maxima, broader thermal optima, and faster rates of ATP hydrolysis and Rubisco reactivation than the predominant Rca isoforms present in cowpea leaves. We showed that a 5-day 38/28 °C heatwave in the absence of water deficit caused leaf temperature to increase to 32 °C. Since this temperature is below the temperature at which Rca activity decreased below 70% of maximum for each Rca isoform, the impact of such a heatwave on Rubisco function and biomass production was only mild. However, cowpea plants growing in smallholder farmer fields in sub-Saharan Africa are likely to experience water deficit alongside warmer heatwaves (Almazroui *et al*., 2020; Song *et al*., 2023). This combination can impact the ability of the plant to cool its leaves leading to higher leaf temperatures (Carmo-Silva *et al*., 2012; Fahad *et al*., 2017; Sato *et al*., 2024). Increasing the abundance of the superior Rca10α and Rca10β isoforms characterised here provides an exciting opportunity to enhance the resilience of cowpea and other crops to future extreme climates.

## Acknowledgements

We thank Dr Sam Taylor (Lancaster University) for input and discussions around experimental design and calculations for determining T_crit_. Ben Davies (Lancaster University) supported plant maintenance, measurements and sampling. We thank Dr Rachel Baxter (Lancaster University) for assistance with data analysis, and Dr Joana Amaral for useful discussions on data interpretation.

## Author contributions

E.C.S conceived and supervised the research. A.G., R.P., I.R., D.J.O., E.C.S. designed experiments. M.T.P. designed and executed the cloning strategy. D.B., P.D.G., D.W. produced and purified recombinant proteins. D.W. and C.J.A. purified Rubisco. D.B. and A.G. performed ATP hydrolysis measurements. A.G. performed Rubisco reactivation measurements. R.P. and I.R. performed heatwave experiments. R.P. performed RNAseq, RT-qPCR and plant growth measurements. R.P. and P.D.G. performed promoter analyses. I.R. performed fluorescence measurements. A.G., R.P. and C.J.A. determined activity and amount of Rubisco and Rubisco activase in leaf extracts. A.G., R.P., C.J.A. and I.R. analysed the data. A.G. and E.C.S. wrote the manuscript with help from all authors.

## Data availability

Data presented are available in the accompanying supporting information and source data. The RNA-seq data reported in this article has been deposited in DDBJ’s Sequence Read Archive (DRA) and are accessible through DRR BioProject accession number PRJDB19806. Processed RNA-seq data are submitted in the DDBJ’s Genomic Expression Archive (GEA) and are accessible through the Experiment accession number (E-GEAD-889). Other data associated with the study are available from the corresponding author.

