## Supporting Information for "Two cowpea Rubisco activase isoforms for crop thermotolerance"

Supporting Information Figures ... 3

Fig. S1. Amino acid residue alignment of the four cowpea Rca isoforms. ... 3

Fig. S2. *In vitro* Rubisco reactivation by the four cowpea Rca isoforms. ... 4

Fig. S3. Experimental design and cabinet temperature monitoring. ... 5

Fig. S4. Cabinet plant layout design. ... 6

Fig. S5. Leaf temperature monitoring over the heatwave experiment. ... 7

Fig. S6. Growth parameters and chlorophyll content during and after heat stress. ... 8

Fig. S7. Method to calculate $T_{crit}$ from chlorophyll fluorescence derived Fv/Fm over a temperature range. ... 9

Fig. S8. Specific Rubisco activities of control and heat-treated plants. ... 10

Fig. S9. Differential gene expression in heat-treated versus control cowpea plants. ... 11

Fig. S10. Volcano plots depicting differential gene expression based on log change across the days of heatwave. ... 12

Fig. S11. Gene expression of heat shock protein 20 (*HSP20*) and the four cowpea *Rca* transcripts. RT-qPCR ... 13

Fig. S12. Identification of potential cis-acting regulatory elements in cowpea Rca promoter regions. ... 14

Fig. S13. Protein abundance of Rca isoforms in leaves of control and heat-treated plants ... 15

Fig. S14. Protocol outline for determination of Rca activity in leaf extracts (LE). ... 16

Fig. S15. Rubisco reactivation by the pool of cowpea Rca isoforms in leaf extracts (LE). ... 17

Supporting Information Tables ... 18

Table S1. Reference protein sequences used to identify cowpea Rca genes. ... 18

Table S2. Primer sequences for adding Golden Gate overhangs to Rca coding regions ... 19

Table S3. Optimum temperature of *in vitro* Rubisco activase (Rca) activity. ... 20

Table S4. Modelling of the *in vitro* temperature response of ATP hydrolysis and Rubisco activation by cowpea Rca isoforms. ... 21

Table S5. RNA sample QC analysis prior to RNA sequencing. ... 22

### Supporting Information Figures

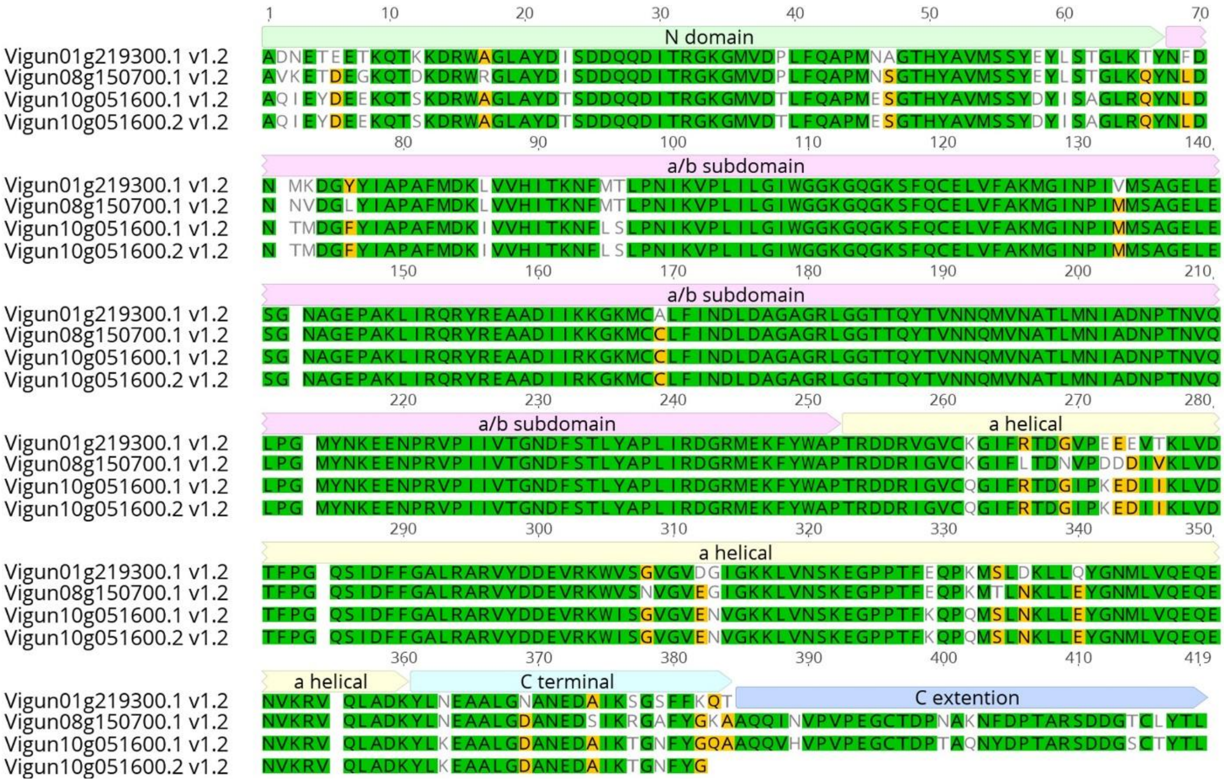

**Fig. S1. Amino acid residue alignment of the four cowpea Rca isoforms.** Shown are mature protein sequences, excluding transit peptides. Rca1β (Vigun01g219300.1 v1.2), Rca8α (Vigun08g150700.1 v1.2), Rca10α (Vigun10g051600.1 v1.2), Rca10β (Vigun10g051600.2 v1.2). Annotations are based on structural domains of *N. tabacum* Rca (Stotz et al., 2011). Sequences were retrieved from Phytozome 13 and the alignment was generated in Geneious 9.1.8.

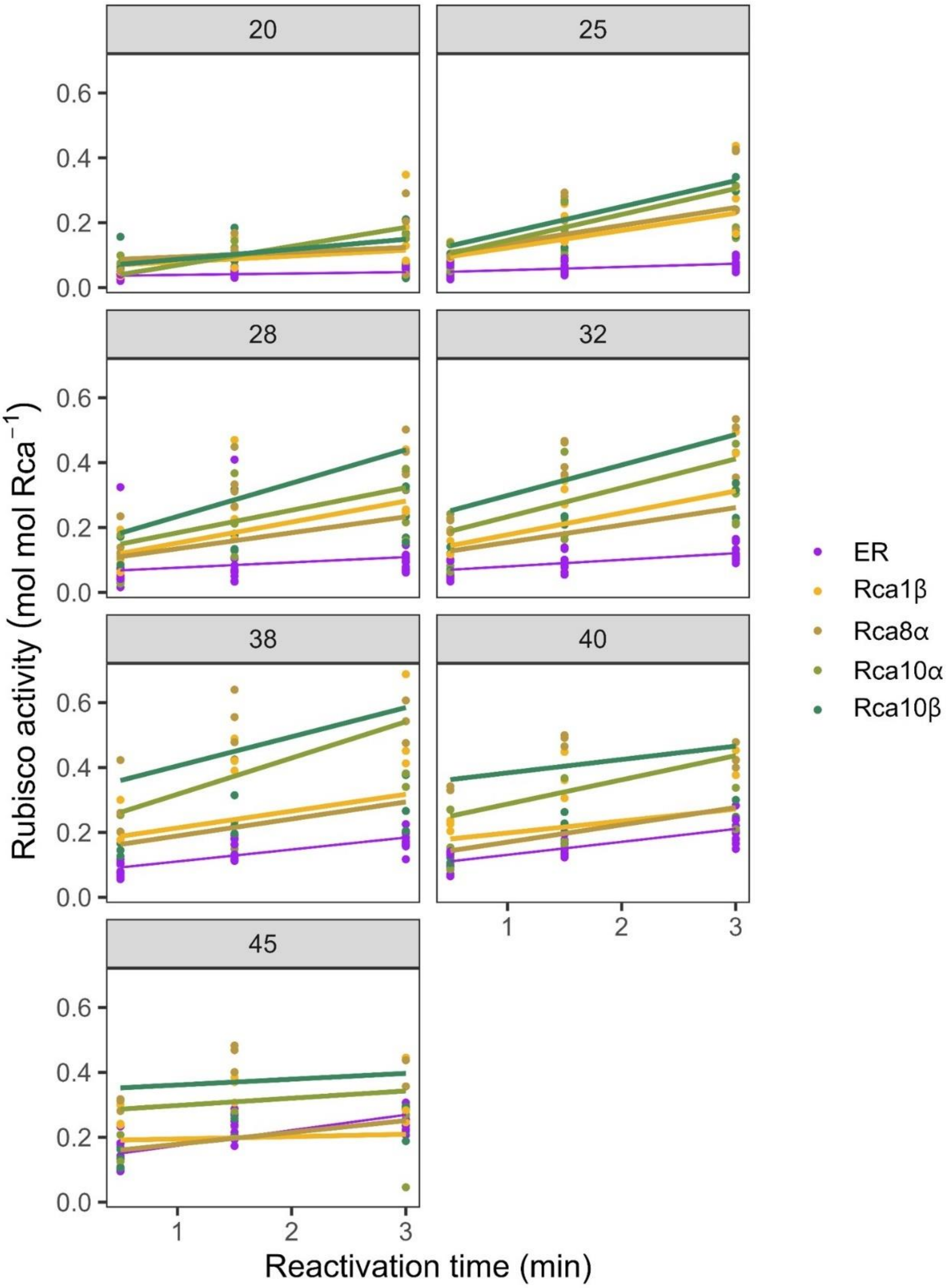

**Fig. S2.** ***In vitro*** **Rubisco reactivation by the four cowpea Rca isoforms.** a) Spontaneous (ER) and Rca mediated reactivation of Rubisco at different timepoints of temperature incubation of the four isoforms Rca1β, Rca8α, Rca10α, Rca10β. For each temperature and each timepoint four technical replicates of ER were performed and one of each Rca isoform. In total for each isoform, activity was determined in three biological replicates (unique purifications) for temperatures 20-32℃ and four for temperatures 38-45℃ (ER n=18, Rca n=3-4). Full sample data provided in source data file.

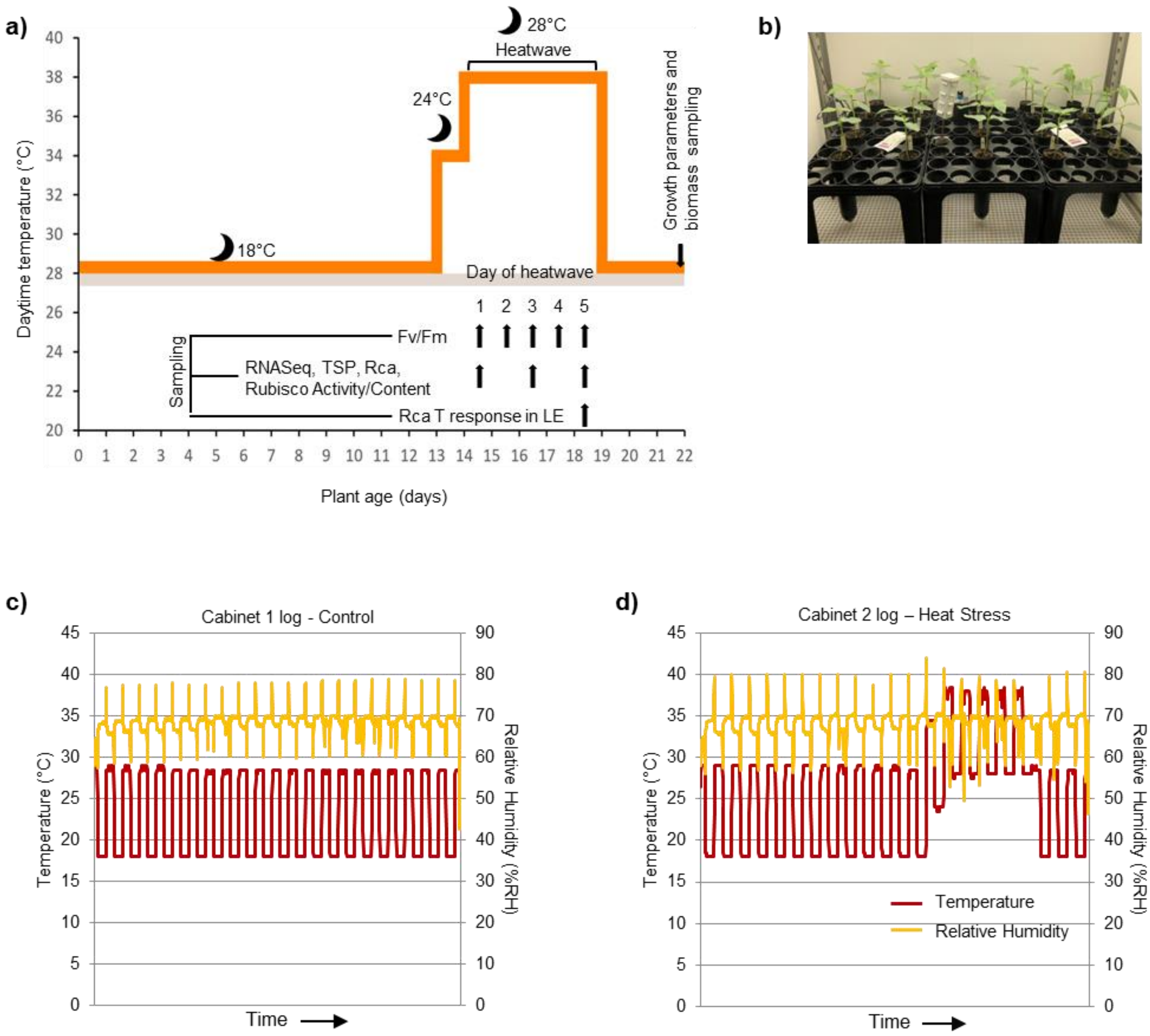

**Fig. S3. Experimental design and cabinet temperature monitoring.** a) Overview of the treatments applied to the plants, where orange = temperatures of heat-treated plants, and grey = temperature of control plants. Night temperatures for each part of the study are shown next to moon symbols. Time of sampling for various analyses is shown by black arrows. b) Picture of plants setup and data logger position. Environmental conditions, air temperature and relative humidity, in a representative control cabinet (c) and heat stress cabinet (d).

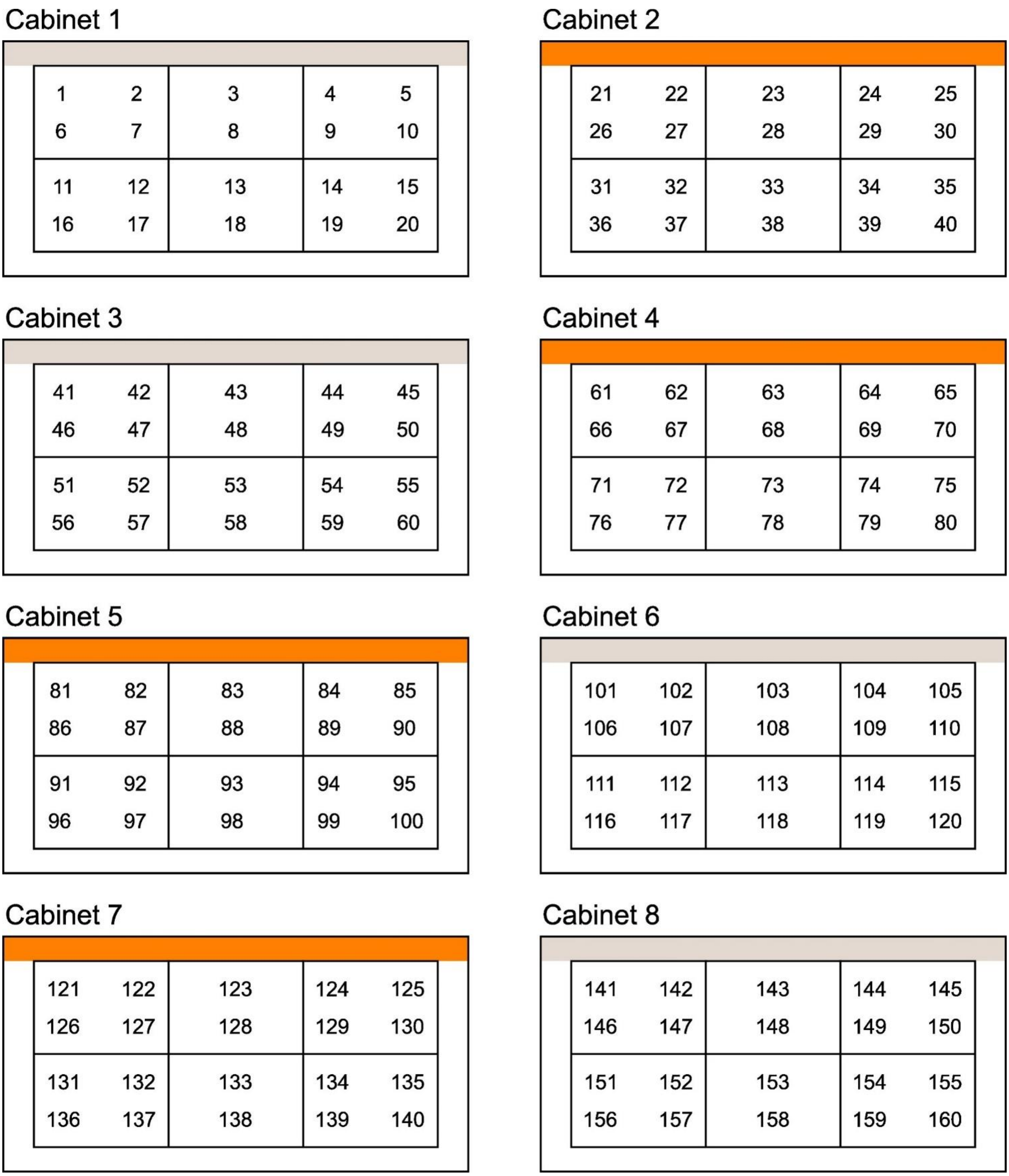

**Fig. S4. Cabinet plant layout design.** Plants were distributed among eight cabinets: four maintained at control temperatures (grey) throughout the experiment and four subjected to a heatwave (orange).

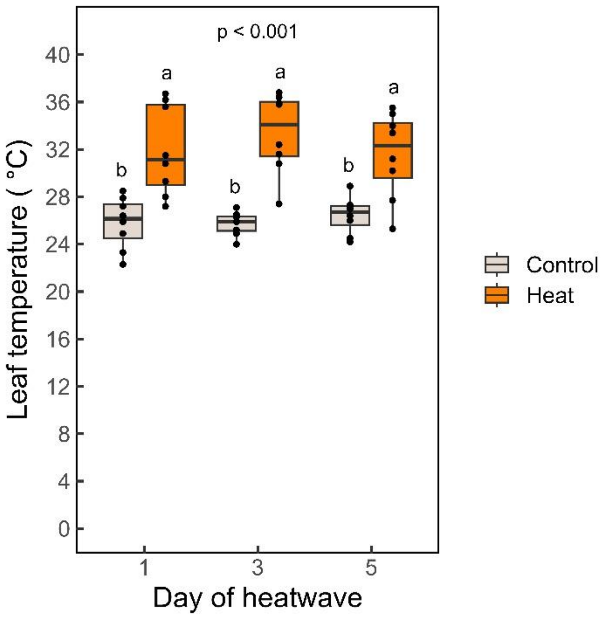

**Fig. S5. Leaf temperature monitoring over the heatwave experiment.** Leaf temperature was measured with a thermal camera on days 1, 3 and 5 of the heatwave, just before taking leaf samples for analysis. There was no significant interaction between days and treatment (p=0.435), the treatment p-value shown was determined using a two-way ANOVA followed by Tukey's post-hoc test (n= 8). The mean leaf temperature across all days for control plants was 26 ± 0.3 °C and for heat-treated plants 32 ± 0.7 °C. Full sample data provided in source data file.

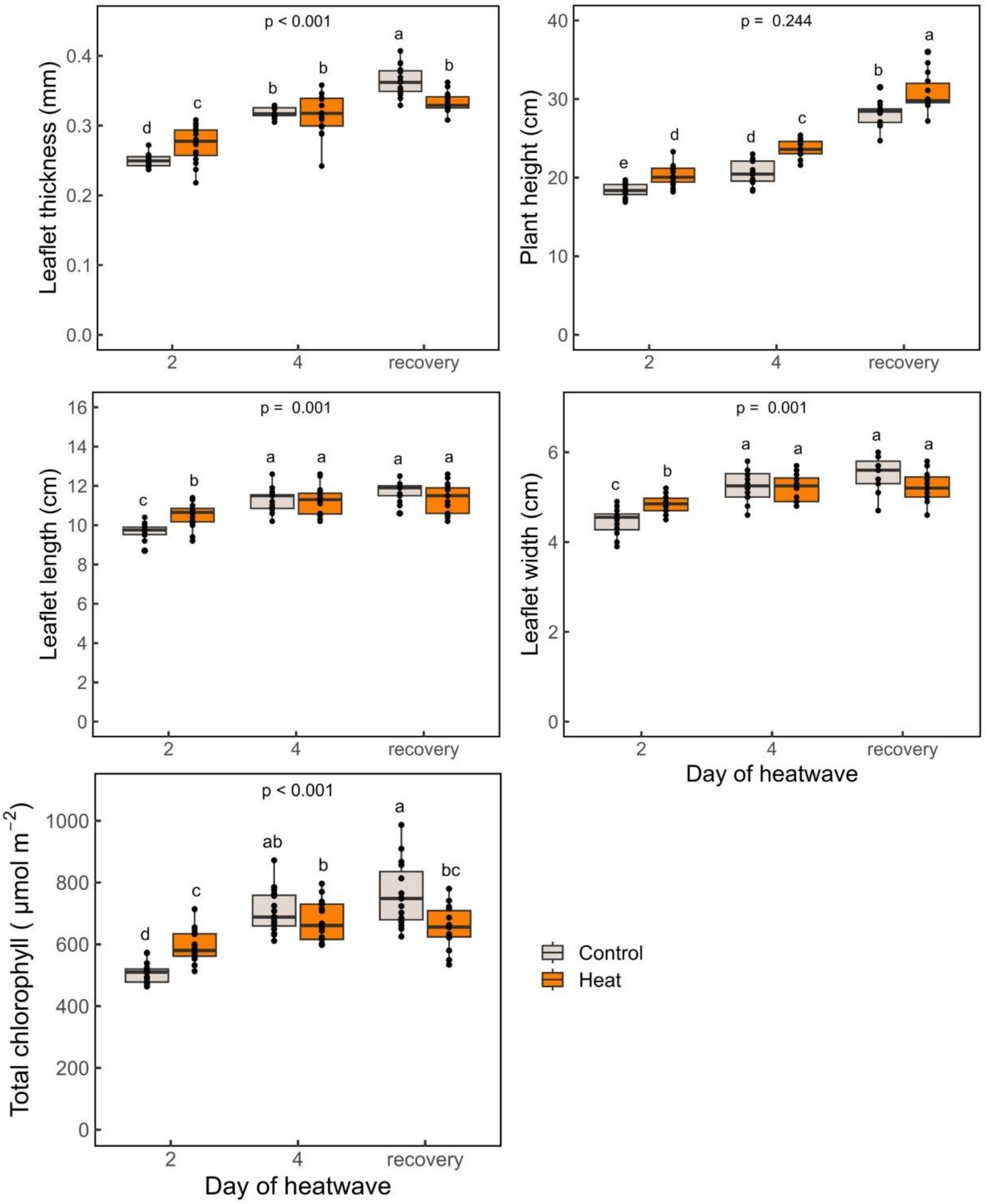

**Fig. S6. Growth parameters and chlorophyll content during and after heat stress.** Chlorophyll content was measured using a handheld meter. Interaction p-values were determined using a two-way ANOVA followed by Tukey's post-hoc test (n=15-16). Full sample data provided in source data file.

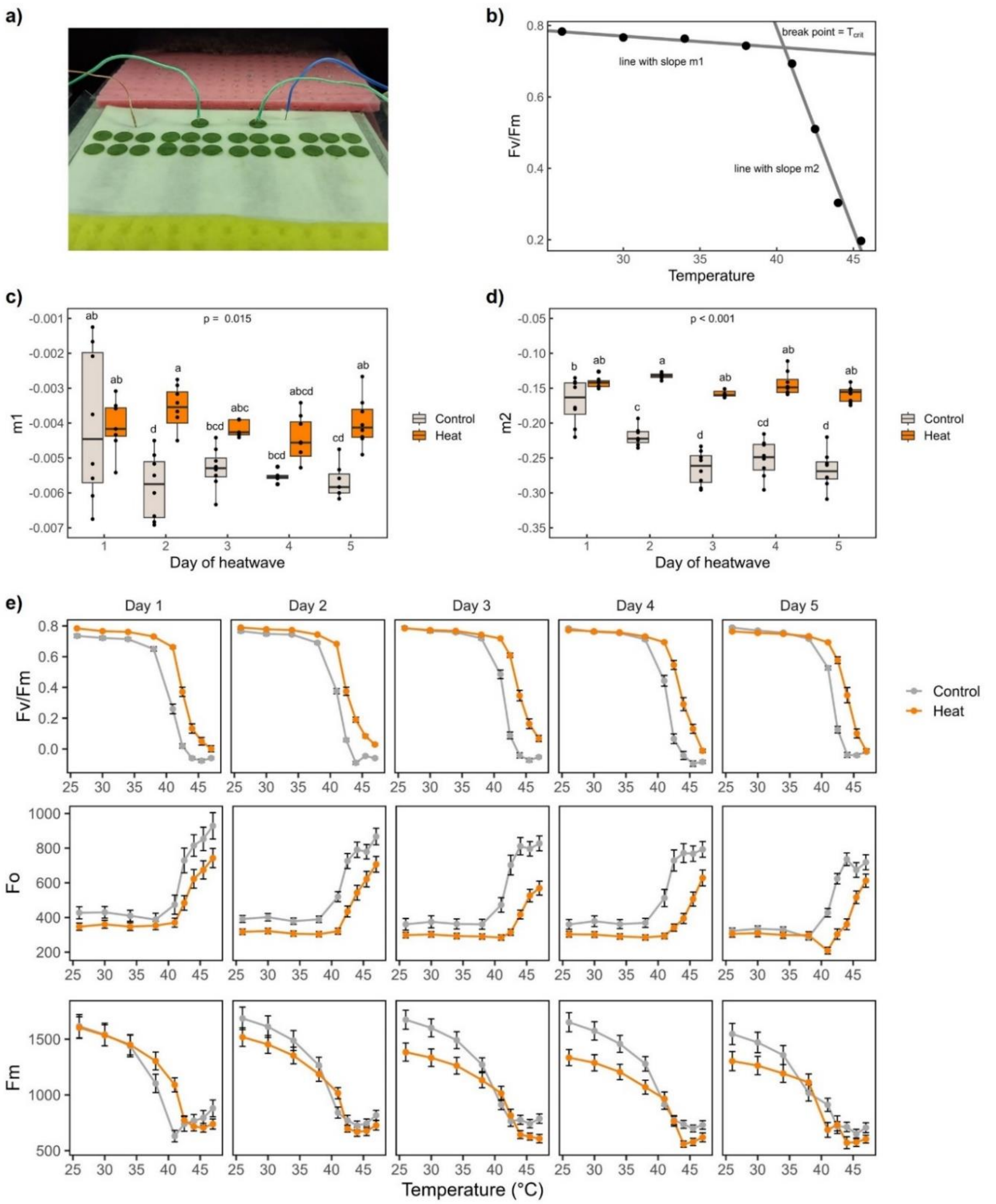

**Fig. S7. Method to calculate $T_{crit}$ from chlorophyll fluorescence derived Fv/Fm over a temperature range.** a) Layout of leaf discs in fluorescence imager showing sample surface with wet filter paper, sponges to provide reservoir of water and thermocouple placement. b) Single example of Fv/Fm values over a temperature range showing the fitted straight lines with slopes $m_1$ & $m_2$ and the breakpoint $T_{crit}$. c) $m_1$ (n=6-7). d) $m_2$ (n=7-8). Interaction p-values were determined using a two-way ANOVA followed by Tukey's post-hoc test. e) Fv/Fm, Fo and Fm over the temperature range and across the five days of heatwave. Points show the treatment averages and error bars show standard errors (n=8). Full sample data provided in source data file.

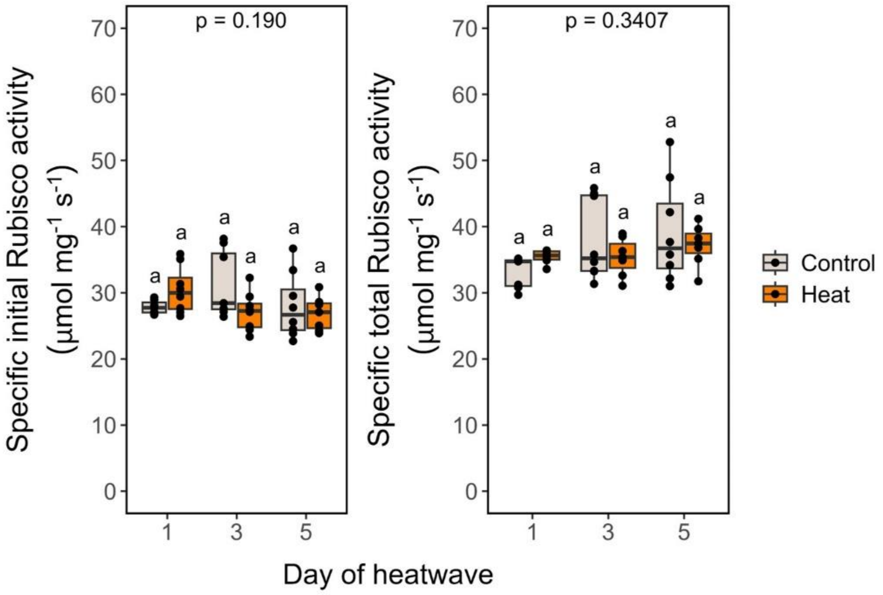

**Fig. S8. Specific Rubisco activities of control and heat-treated plants.** Initial and total activities were normalised to the Rubisco concentration. Interaction p-values were determined using a two-way ANOVA followed by Tukey's post-hoc test (n=6-8). Full sample data are included in the source data file.

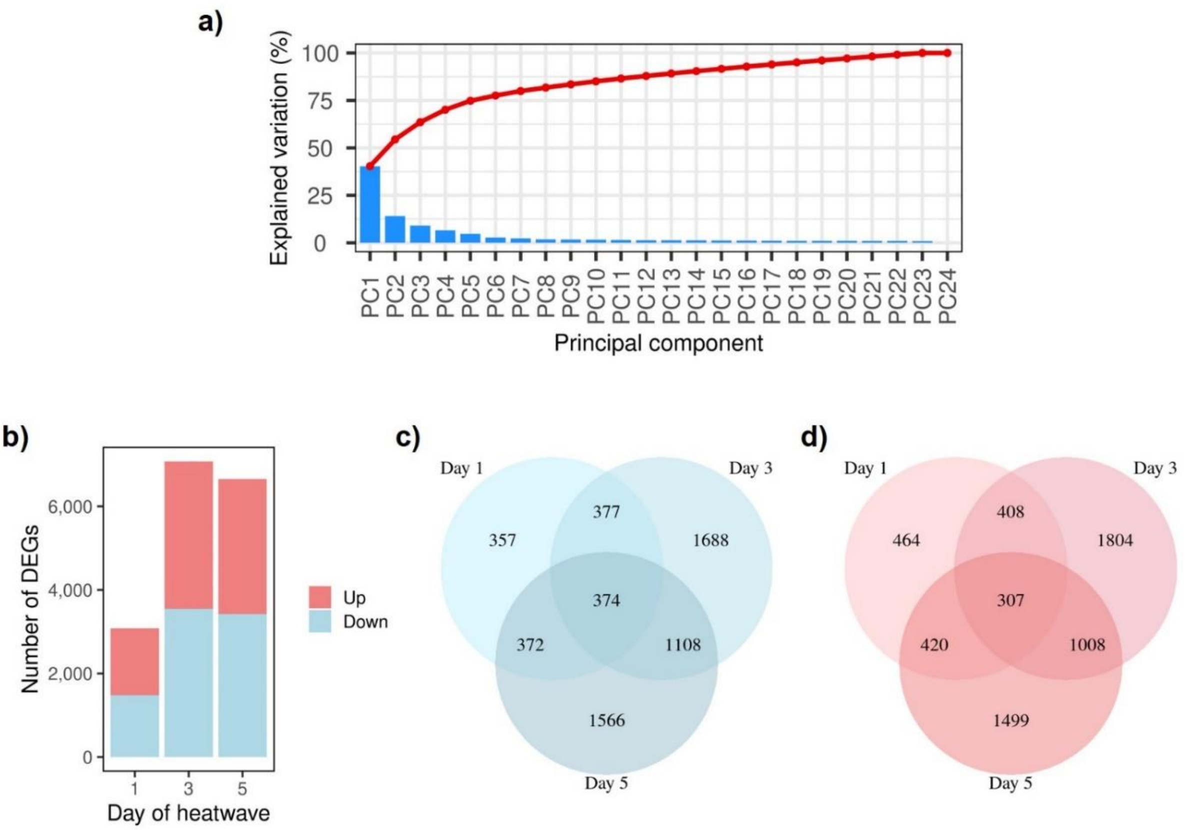

**Fig. S9. Differential gene expression in heat-treated versus control cowpea plants.** Young plants were exposed to a +10 °C heatwave for 5 days and samples taken for RNA-seq analysis on days 1, 3, 5. a) Scree plot showing the principal components (PCA) driving differentially expressed genes (DEGs) in control and heat-treated plants. b) The number of DEGs that are downregulated and upregulated at days 1, 3, and 5 of the heat treatment. (c, d) Venn diagrams showing the number of unique and common downregulated (c) and upregulated (d) DEGs in heat-treated compared to control plants on days 1, 3 and 5. Venn diagrams were produced using the VennDiagram in R studio.

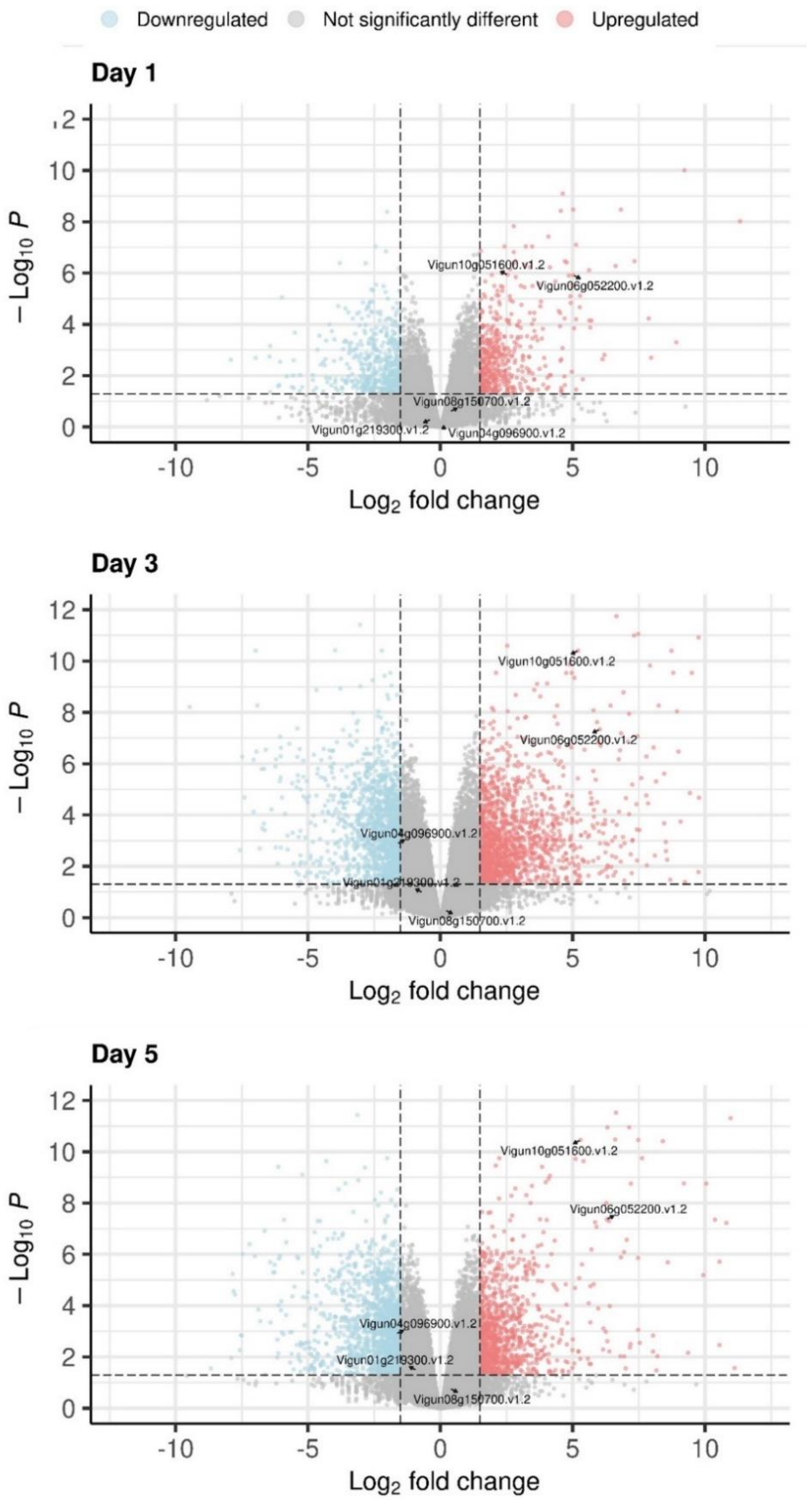

**Fig. S10. Volcano plots depicting differential gene expression based on log change across the days of heatwave.** Blue corresponds to the downregulated genes, coral red to upregulated gene expression with positive log fold change during the heatwave while middle grey corresponds to the genes whose expression was unaffected by the heat treatment. Labels correspond to the *HSP20* (*Vigun06g052200*.v1.2), *RbcS* (*Vigun04g096900*.v1.2) and *Rca* encoding genes (*Rca1*: *Vigun01g219300*.v1.2, *Rca8*: *Vigun08g150700*.v1.2, *Rca10*: *Vigun10g051600*.v1.2).

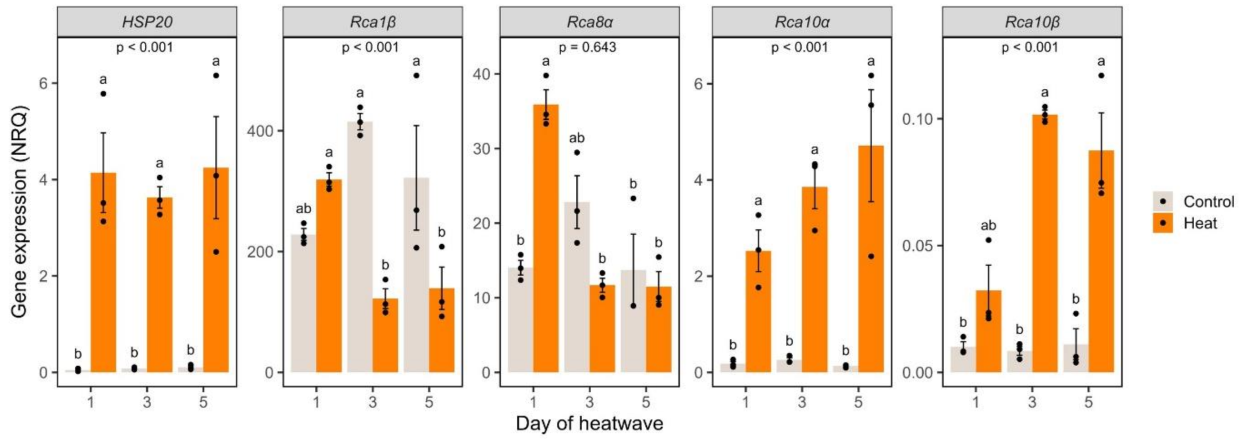

**Fig. S11. Gene expression of heat shock protein 20 (*HSP20*) and the four cowpea *Rca* transcripts.** RT-qPCR was performed on samples taken at days 1,3, and 5 of the heatwave, in a replicate experiment using the same design as the RNA-seq analyses (Fig. 2a). RNA extraction and RT-qPCR analysis information is provided in the MIQE checklist (Table S7) and primer sequences and information in Supplementary Table 8. Key results are confirmatory of the RNA-seq analysis, showing a significant increase in gene expression of *Rca10α* and *Rca10β* during the heatwave. Two-way ANOVA followed by Tukey's post-hoc test was performed after log transformation (n=3). P-values correspond to heat treatment effect across the days of the heatwave for each of the genes. Full sample data provided in source data file.

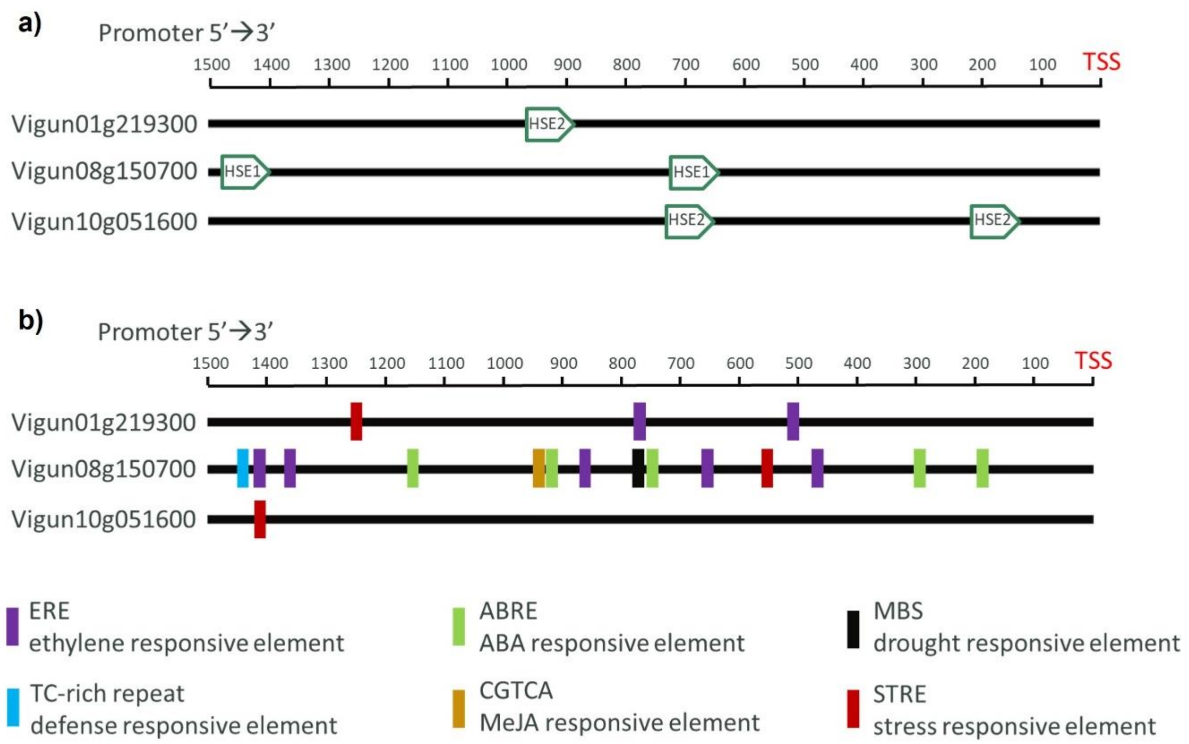

c)

| Cis Regulatory Motif | VuRca1 | VuRca8 | VuRca10 | Signal Seq | Function |
|---|---|---|---|---|---|
| TATABOXOSPAL | | 2 | 3 | TATTTAA | BIOSYNTHESIS, DEFENCE, DROUGHT STRESS |
| WBOXNTERF3 | 1 | 2 | 3 | TGACY | DEFENCE, STRESS |
| GT1GMSCAM4 | 6 | 6 | 6 | GAAAAA | DEFENCE, STRESS |
| PRECONSCRHSP70A | 1 | | | SCGAYNRNNNNNNNNNNNNNNNHD | LIGHT/ STRESS? |
| MYB1AT | 4 | 3 | | WAACCA | STRESS |
| MYCCONSENSUSAT | 6 | 14 | | CANNTG | STRESS |
| MYB2CONSENSUSAT | | 2 | | YAACKG | STRESS |
| BIHD1OS | 2 | 8 | 2 | TGTCA | STRESS |
| TATABOX3 | 2 | 2 | 2 | TATTAAT | STRESS |
| MYBCORE | 1 | 2 | | CNGTTR | STRESS |
| ABREZMRAB28 | 2 | | | CCACGTGG | STRESS |
| EMBP1TAEM | 1 | | | CACGTGGC | STRESS |
| IRO2OS | 2 | | | CACGTGG | STRESS |
| ABREATCONSENSUS | 1 | | | YACGTGGC | STRESS |
| ACGTATERD1 | 4 | 8 | 2 | ACGT | STRESS |
| MYB2AT | | 1 | | TAACTG | STRESS |
| MYCATERD1 | 1 | 3 | | CATGTG | STRESS |
| MYCATRD22 | 1 | 3 | | CACATG | STRESS |
| CURECORECR | 2 | 2 | 4 | GTAC | STRESS, COPPER DEFICIENCY |
| CCAATBOX1 | 8 | 5 | 2 | CCAAT | STRESS, HEAT SHOCK, TEMPERATURE |
| ABRERATCAL | 2 | 2 | | MACGYGB | STRESS, HORMONE |
| ACGTABREMOTIFA2OSEM | 1 | 2 | | ACGTGKC | STRESS, HORMONE |
| ABRELATERD1 | 1 | 5 | | ACGTG | STRESS, HORMONE |
| SURECOREATSULTR11 | 5 | | 3 | GAGAC | SULPHUR DEFICIENCY, STRESS, HORMONE |
| LTRECOREATCOR15 | 1 | | | CCGAC | TEMP, LIGHT |
| CBFHV | | | 2 | RYCGAC | TEMPERATURE |
| CRTDREHVCBF2 | | | 2 | GTCGAC | TEMPERATURE |

**Fig. S12. Identification of potential cis-acting regulatory elements in cowpea Rca promoter regions.** a) To identify potential cis-acting regulatory elements in cowpea *Rca* promoter regions, 1.5kb regions upstream of the transcriptional start sites (TSSs) were investigated for heat stress elements (HSE) based on consensus sequences identified by Jung *et al.* (2013). Several instances of consensus HSE sequences HSE1 (GAAnnTTC) and HSE2 (TTCnnGAA) were identified. The same promoter regions were analysed using PlantCARE (b) and PLACE (c) to determine the number of motifs related to temperature and stress response.

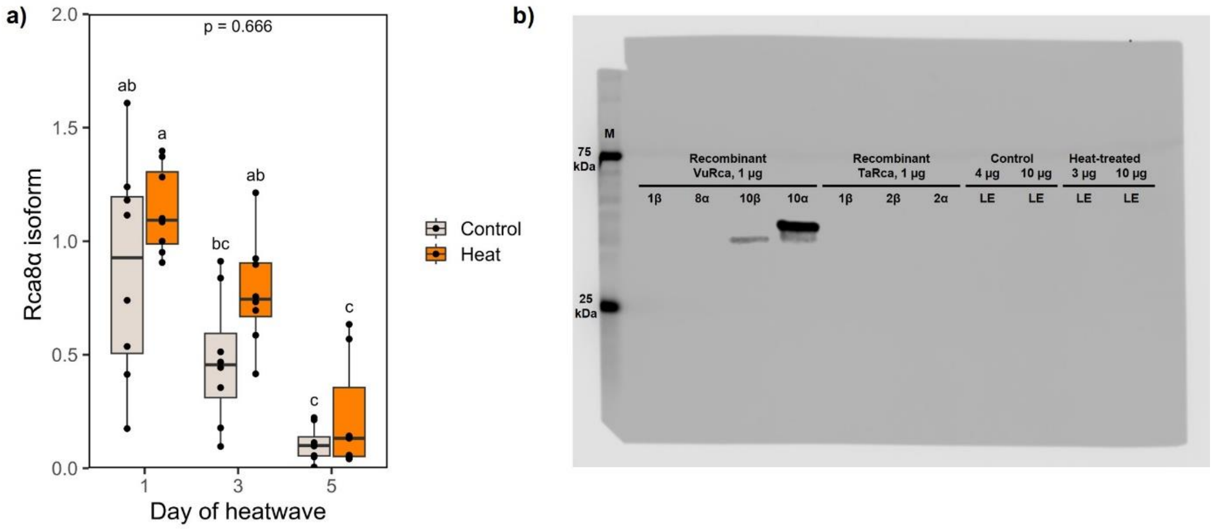

**Fig. S13. Protein abundance of Rca isoforms in leaves of control and heat-treated plants**. a) A specific antibody that reacts only with Rca8α (Bloemers & Carmo-Silva, 2024) was used to quantify the abundance of this protein isoform. Anti-Rca8α antibody was generated using phage display where the selected peptide targeted residues 376-391 of the mature protein sequence (KRGAFYGKAAQQINVP)(Bloemers & Carmo-Silva, 2024). Blots were analysed as described in Methods. Interaction p-values were calculated using two-way ANOVA followed by Tukey post-hoc test (n=7-8). Full sample data are provided in the source data file. b) An anti-Rca10α/β antibody was also generated using phage display, where the selected peptide targeted residues 376-391 of the mature protein sequence (KTGNFYGQAAQQVHVP). The anti-Rca10α/β antibody showed detection of recombinant VuRca10, however, no visible bands were seen for control and heat-treated leaf extract (LE) samples. Both the leaf extract samples tested were sampled on day 5 of the heatwave. There was also no binding of the anti-Rca10α/β antibody to recombinant wheat Rca (TaRca).

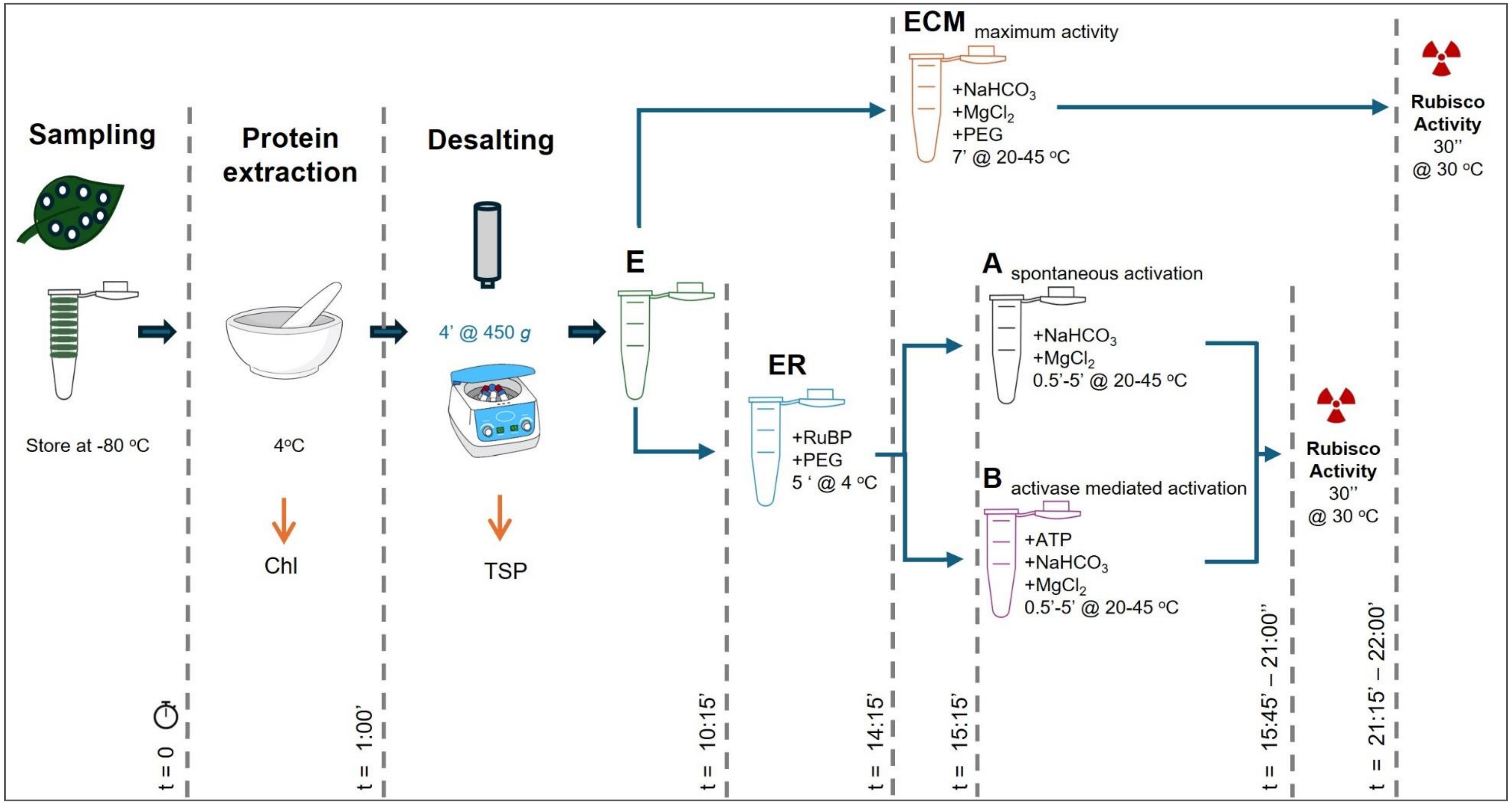

**Fig. S14. Protocol outline for determination of Rca activity in leaf extracts (LE).** Samples consisting of eight leaf discs per plant were collected on day 5 of the heatwave. The protocol timeline starts (t=0) with protein extraction by grinding leaves in ice-cold extraction buffer. Homogenate aliquots were used for leaf chlorophyll content determination. A spin desalting step enables removal of $Mg^{2+}$ ions and $CO_2$ to obtain free uncarbarmylated Rubisco (E). Aliquots of the desalted leaf extract were taken for total soluble protein (TSP), activity of fully carbamylated Rubisco (ECM) and inhibition of uncarbamylated Rubisco by binding to RuBP (ER) for Rca activity determination. The ECM sample was incubated with 5% PEG3350, 10 mM $NaHCO_3$, and 30 mM $MgCl_2$ for 7 min at 20-45 °C prior to measuring Rubisco activity at 30 °C. The ER sample was supplemented with 5% PEG3350 and 4 mM RuBP and incubated for 5 min at 4 °C to form the inhibited Rubisco–RuBP (ER) complex. The leaf extract containing ER and Rca was then used to initiate reactivation assays with 10 mM $NaHCO_3$ and 30 mM $MgCl_2$ in the presence or absence of 5 mM ATP plus an ATP-regenerating system at 20-45 °C to determine spontaneous (A) and activase-mediated (B) activation of Rubisco by measuring the increase in Rubisco activity at 30 °C.

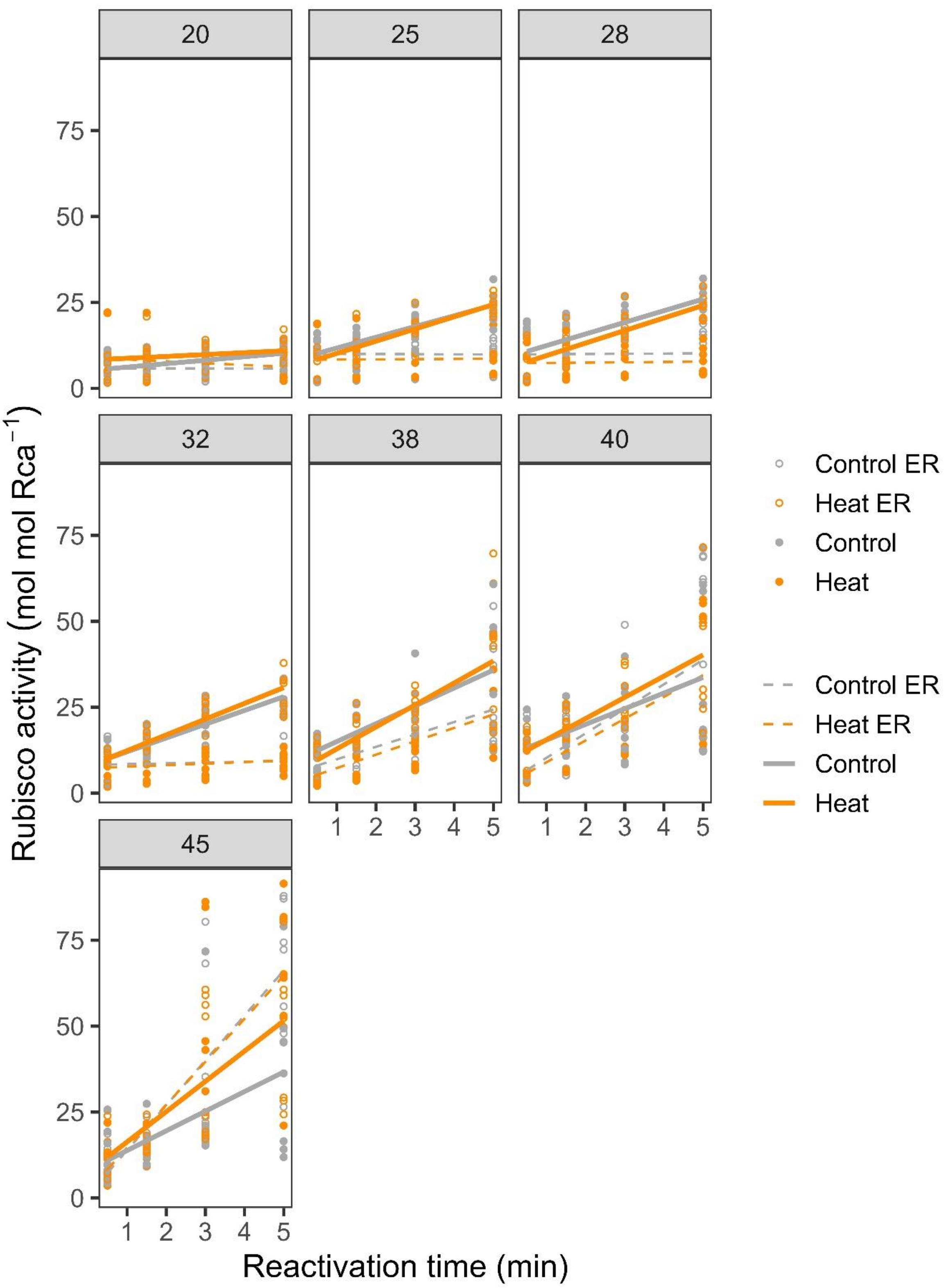

**Fig. S15. Rubisco reactivation by the pool of cowpea Rca isoforms in leaf extracts (LE).** Spontaneous (open circles, dashed line) and Rca mediated (filled circles, solid line) reactivation of Rubisco at different timepoints of temperature incubation of control (grey) and heat-treated (orange) plants (n=7-8). Samples were collected on day 5 of the heatwave. Symbols represent individual measurements. Full sample data provided in source data file.

### Supporting Information Tables

**Table S1. Reference protein sequences used to identify cowpea Rca genes.**

| Species | Protein Name | Identifier (Database) |
|---|---|---|
| *Arabidopsis thaliana* | AtRca | At2g39730 (TAIR) |
| *Nicotiana tabacum* | NtRca | Q40460 (Uniprot) |
| *Triticum aestivum* | TaRca1 | TraesCS4A02G177600 (EnsemblPlants) |
| | TaRca2 | TraesCS4A02G177500 (EnsemblPlants) |

**Table S2. Primer sequences for adding Golden Gate overhangs to Rca coding regions**. For each Rca coding sequence a specific primer pair was used, except for Rca10β where the forward primer for Rca10α (Rca10.1A_GG_F) could be used due to the identical coding sequence at the 5'-end.

| Primer Name | Sequence (5' > 3') | Notes |
|---|---|---|
| Rca1.1B_GG_F | GCGGTCTCACAATGCCGACAACGAAACGGAG | Adds Golden Gate sites to Rca1 |
| Rca1.1B_GG_R | CAGGTCTCGATCATTAAGTCTGTTTGAAGAAGC | |
| Rca8.1A_GG_F | GCGGTCTCACAATGCAGTCAAAGAAACCGATG | Adds Golden Gate sites to Rca8 |
| Rca8.1A_GG_R | CAGGTCTCGATCATCACAGAGTGTACAGGCAC | |
| Rca10.1A_GG_F | GCGGTCTCACAATGCGCAGATCGAATATGACG | Adds Golden Gate sites to Rca10a |
| Rca10.1A_GG_R | CAGGTCTCGATCACTACAGGGTGTACGTGCAG | |
| Rca10.2B_GG_R | CAGGTCTCGATCATCAACCGTAGAAGTTACCGGT | Adds Golden Gate site to Rca10b |

**Table S3. Optimum temperature of *in vitro* Rubisco activase (Rca) activity.** Maximum rates of ATP hydrolysis and Rubisco reactivation, and corresponding temperature of maximum activity ($T_{max}$), optimum temperature range ($T_{opt}$, above 70% activity) and temperature above the optimum at which 50% of the maximum activity remains ($T_{0.5}$), as estimated from the best-fit models applied to describe the *in vitro* temperature response of each Rca isoform (Table S4). The best fitting model was applied to the combined biological replicates presented in Fig. 1b (n=3-4).

| **Rca Isoform** | **$ATPase_{max}$ (mol min$^{-1}$ mol Rca$^{-1}$)** | **$T_{max}$ (°C)** | **$T_{opt}$ (°C)** | **$T_{0.5}$ (°C)** |
|---|---|---|---|---|
| **1β** | 27.6 | 36.5 | 30.3 – 42.5 | 44.5 |
| **8α** | 38.6 | 35.8 | 28.5 – 42.9 | 45.3 |
| **10α** | 176.1 | 41.9 | 30.4 – 51.0 | 53.4 |
| **10β** | 179.3 | 39.3 | 27.8 – 50.7 | 54.0 |

| **Rca Isoform** | **Rubisco $reactivation_{max}$ (mol $R_{A.S.}$ min$^{-1}$ mol Rca$^{-1}$)** | **$T_{max}$ (°C)** | **$T_{opt}$ (°C)** | **$T_{0.5}$ (°C)** |
|---|---|---|---|---|
| **1β** | 0.09 | 30.7 | 25.8 – 35.4 | 37.0 |
| **8α** | 0.08 | 27.8 | 22.2 – 35.2 | 38.5 |
| **10α** | 0.14 | 33.9 | 25.4 – 40.5 | 42.3 |
| **10β** | 0.19 | 32.6 | 24.3 – 40.9 | 43.0 |

**Table S4. Modelling of the *in vitro* temperature response of ATP hydrolysis and Rubisco activation by cowpea Rca isoforms.** The model providing the best fit to the data is highlighted in bold and was selected according to the lowest AIC score (Akaike information criterion) calculated according to Akaike (1974) using the AIC function in R. Models were applied to the full dataset shown in Fig. 1b (n=3-4).

| Rca Isoform | **ATPase assay** | | | **Rubisco activation assay** | | |
|---|---|---|---|---|---|---|
| | Model | degrees of freedom (K) | AIC score | Model | degrees of freedom (K) | AIC score |
| 1β | $2^{nd}$ order poly | 4 | 217 | $2^{nd}$ order poly | 4 | -76 |
| | $3^{rd}$ order poly | 5 | 218 | $3^{rd}$ order poly | 5 | -76 |
| | **$4^{th}$ order poly** | **6** | **210** | **$4^{th}$ order poly** | **6** | **-77** |
| | $5^{th}$ order poly | 7 | 212 | $5^{th}$ order poly | 7 | -75 |
| | GAM | 3 | 236 | GAM | 3 | -72 |
| 8α | $2^{nd}$ order poly | 4 | 177 | $2^{nd}$ order poly | 4 | -82.6 |
| | $3^{rd}$ order poly | 5 | 178 | **$3^{rd}$ order poly** | **5** | **-82.95** |
| | **$4^{th}$ order poly** | **6** | **174** | $4^{th}$ order poly | 6 | -81 |
| | $5^{th}$ order poly | 7 | 176 | $5^{th}$ order poly | 7 | -79.9 |
| | GAM | 3 | 194 | GAM | 3 | -80 |
| 10α | $2^{nd}$ order poly | 4 | 350 | $2^{nd}$ order poly | 4 | -84.59 |
| | **$3^{rd}$ order poly** | **5** | **349** | **$3^{rd}$ order poly** | **5** | **-84.61** |
| | $4^{th}$ order poly | 6 | 351 | $4^{th}$ order poly | 6 | -83 |
| | $5^{th}$ order poly | 7 | 352 | $5^{th}$ order poly | 7 | -82 |
| | GAM | 3 | 365 | GAM | 3 | -67 |
| 10β | **$2^{nd}$ order poly** | **4** | **368** | **$2^{nd}$ order poly** | **4** | **-86** |
| | $3^{rd}$ order poly | 5 | 369 | $3^{rd}$ order poly | **5** | -84.5 |
| | $4^{th}$ order poly | 6 | 371 | $4^{th}$ order poly | 6 | -83 |
| | $5^{th}$ order poly | 7 | 372 | $5^{th}$ order poly | 7 | -82.9 |
| | GAM | 3 | 375 | GAM | 3 | -61 |

**Table S5. RNA sample QC analysis prior to RNA sequencing.**

| | | | MICROPLATE READER | | | NOVOGENE | | | |
|---|---|---|---|---|---|---|---|---|---|
| **Sample ID** | **Treatment** | **Timepoint** | **Concentration (ng/µl)** | **260/280 ratio** | **260/230 ratio** | **Concentration (ng/µl)** | **Volume (µl)** | **Total amount (µg)** | **Integrity value (RIN)** |
| RP001 | Control | D5 | 311.75 | 2.01 | 2.31 | 236.46 | 41 | 9.69 | 7.1 |
| RP002 | Control | D1 | 284.21 | 2.01 | 2.40 | 228.61 | 42 | 9.60 | 8.3 |
| RP020 | Control | D3 | 725.36 | 2.05 | 2.39 | 866.62 | 42 | 36.40 | 8.8 |
| RP031 | Heat | D1 | 488.38 | 2.02 | 2.38 | 529.17 | 42 | 22.23 | 8.5 |
| RP038 | Heat | D5 | 80.82 | 1.98 | 2.29 | 50.29 | 42 | 2.11 | 7.7 |
| RP040 | Heat | D3 | 103.86 | 1.99 | 2.16 | 70.71 | 42 | 2.97 | 8.0 |
| RP051 | Control | D5 | 321.12 | 2.01 | 2.40 | 240.84 | 42 | 10.12 | 7.4 |
| RP058 | Control | D1 | 481.18 | 2.03 | 2.38 | 559.28 | 42 | 23.49 | 9.0 |
| RP059 | Control | D3 | 340.80 | 2.02 | 2.27 | 282.79 | 38 | 10.75 | 7.6 |
| RP070 | Heat | D5 | 121.95 | 2.00 | 2.19 | 86.02 | 42 | 3.61 | 7.3 |
| RP077 | Heat | D1 | 785.70 | 2.05 | 2.39 | 1005.5 | 42 | 42.23 | 8.5 |
| RP078 | Heat | D3 | 192.73 | 2.00 | 2.33 | 130.67 | 42 | 5.49 | 7.9 |
| RP086 | Heat | D5 | 144.70 | 1.98 | 2.22 | 99.35 | 42 | 4.17 | 7.7 |
| RP092 | Heat | D1 | 582.69 | 2.03 | 2.39 | 702.46 | 42 | 29.50 | 8.9 |
| RP098 | Heat | D3 | 293.68 | 2.01 | 2.37 | 226.71 | 42 | 9.52 | 7.8 |
| RP111 | Control | D3 | 519.96 | 2.03 | 2.32 | 612.69 | 42 | 25.73 | 8.8 |
| RP115 | Control | D5 | 144.46 | 2.00 | 2.29 | 105.08 | 38 | 3.99 | 6.8 |
| RP120 | Control | D1 | 458.21 | 2.02 | 2.38 | 499.47 | 41 | 20.48 | 8.9 |
| RP121 | Heat | D5 | 133.62 | 1.99 | 2.24 | 86.53 | 43 | 3.72 | 6.3 |
| RP122 | Heat | D3 | 236.64 | 2.00 | 2.38 | 171.98 | 42 | 7.22 | 8.2 |
| RP130 | Heat | D1 | 764.49 | 2.05 | 2.40 | 977.39 | 42 | 41.05 | 8.7 |
| RP141 | Control | D3 | 600.24 | 2.04 | 2.40 | 659.07 | 41 | 27.02 | 8.6 |
| RP144 | Control | D5 | 137.09 | 2.00 | 2.24 | 93.48 | 40 | 3.74 | 7.7 |
| RP159 | Control | D1 | 698.14 | 2.05 | 2.40 | 802.96 | 40 | 32.12 | 8.9 |

#### Table S6. Sequencing and alignment statistics.

| Sample ID | Treatment | Timepoint | Data_file_size (bytes) | Number_seq_raw | Number_seq_clean | Number_seq_removed | Percentage_seq_removed | Total_bases_raw (Gbp) | Total_bases_clean (Gbp) | GC_content (%) | Mean_quality_per_read (Phred score) |
|---|---|---|---|---|---|---|---|---|---|---|---|
| RP001_1 | control | D5 | 13425721087 | 36045152 | 35537602 | 507550 | 1.41 | 5.4 | 5.2 | 46 | 36 |
| RP001_2 | control | D5 | 13425721087 | 36045152 | 35537602 | 507550 | 1.41 | 5.4 | 5.2 | 46 | 36 |
| RP002_1 | control | D1 | 11605151548 | 31157275 | 30748411 | 408864 | 1.31 | 4.6 | 4.5 | 46 | 36 |
| RP002_2 | control | D1 | 11605151548 | 31157275 | 30748411 | 408864 | 1.31 | 4.6 | 4.5 | 46 | 36 |
| RP020_1 | control | D3 | 12056176450 | 32368235 | 31981933 | 386302 | 1.19 | 4.8 | 4.7 | 46 | 36 |
| RP020_2 | control | D3 | 12056176450 | 32368235 | 31981933 | 386302 | 1.19 | 4.8 | 4.7 | 46 | 36 |
| RP031_1 | heat | D1 | 13190424154 | 35413441 | 34930680 | 482761 | 1.36 | 5.3 | 5.1 | 46 | 36 |
| RP031_2 | heat | D1 | 13190424154 | 35413441 | 34930680 | 482761 | 1.36 | 5.3 | 5.1 | 46 | 36 |
| RP038_1 | heat | D5 | 12819613510 | 34417863 | 33977748 | 440115 | 1.28 | 5.1 | 5.0 | 44 | 36 |
| RP038_2 | heat | D5 | 12819613510 | 34417863 | 33977748 | 440115 | 1.28 | 5.1 | 5.0 | 44 | 36 |
| RP040_1 | heat | D3 | 14684686925 | 39425215 | 38884497 | 540718 | 1.37 | 5.9 | 5.7 | 45 | 36 |
| RP040_2 | heat | D3 | 14684686925 | 39425215 | 38884497 | 540718 | 1.37 | 5.9 | 5.7 | 45 | 36 |
| RP051_1 | control | D5 | 15268299964 | 40992041 | 40445324 | 546717 | 1.33 | 6.1 | 6.0 | 46 | 36 |
| RP051_2 | control | D5 | 15268299964 | 40992041 | 40445324 | 546717 | 1.33 | 6.1 | 6.0 | 46 | 36 |
| RP058_1 | control | D1 | 17162706419 | 46078190 | 45422671 | 655519 | 1.42 | 6.9 | 6.7 | 46 | 36 |
| RP058_2 | control | D1 | 17162706419 | 46078190 | 45422671 | 655519 | 1.42 | 6.9 | 6.7 | 46 | 36 |
| RP059_1 | control | D3 | 14002141251 | 37592419 | 37121062 | 471357 | 1.25 | 5.6 | 5.5 | 46 | 36 |
| RP059_2 | control | D3 | 14002141251 | 37592419 | 37121062 | 471357 | 1.25 | 5.6 | 5.5 | 46 | 36 |
| RP070_1 | heat | D5 | 13799927700 | 37049814 | 36573118 | 476696 | 1.29 | 5.5 | 5.4 | 44 | 36 |
| RP070_2 | heat | D5 | 13799927700 | 37049814 | 36573118 | 476696 | 1.29 | 5.5 | 5.4 | 44 | 36 |
| RP077_1 | heat | D1 | 12530825305 | 33642540 | 33192110 | 450430 | 1.34 | 5.0 | 4.9 | 46 | 36 |
| RP077_2 | heat | D1 | 12530825305 | 33642540 | 33192110 | 450430 | 1.34 | 5.0 | 4.9 | 46 | 36 |
| RP078_1 | heat | D3 | 11027898150 | 29607379 | 29210364 | 397015 | 1.34 | 4.4 | 4.3 | 45 | 36 |
| RP078_2 | heat | D3 | 11027898150 | 29607379 | 29210364 | 397015 | 1.34 | 4.4 | 4.3 | 45 | 36 |
| RP086_1 | heat | D5 | 11671673556 | 31335849 | 30838307 | 497542 | 1.59 | 4.7 | 4.5 | 45 | 36 |
| RP086_2 | heat | D5 | 11671673556 | 31335849 | 30838307 | 497542 | 1.59 | 4.7 | 4.5 | 45 | 36 |
| RP092_1 | heat | D1 | 12620133920 | 33882356 | 33446049 | 436307 | 1.29 | 5.0 | 4.9 | 46 | 36 |
| RP092_2 | heat | D1 | 12620133920 | 33882356 | 33446049 | 436307 | 1.29 | 5.0 | 4.9 | 46 | 36 |
| RP098_1 | heat | D3 | 11830130671 | 31761357 | 31313377 | 447980 | 1.41 | 4.7 | 4.6 | 45 | 36 |
| RP098_2 | heat | D3 | 11830130671 | 31761357 | 31313377 | 447980 | 1.41 | 4.7 | 4.6 | 45 | 36 |
| RP111_1 | control | D3 | 13624866297 | 36579818 | 36114521 | 465297 | 1.27 | 5.4 | 5.3 | 46 | 36 |
| RP111_2 | control | D3 | 13624866297 | 36579818 | 36114521 | 465297 | 1.27 | 5.4 | 5.3 | 46 | 36 |
| RP115_1 | control | D5 | 13725270653 | 36849316 | 36385324 | 463992 | 1.26 | 5.5 | 5.4 | 45 | 36 |
| RP115_2 | control | D5 | 13725270653 | 36849316 | 36385324 | 463992 | 1.26 | 5.5 | 5.4 | 45 | 36 |
| RP120_1 | control | D1 | 14283643262 | 38348422 | 37812563 | 535859 | 1.40 | 5.7 | 5.6 | 46 | 36 |
| RP120_2 | control | D1 | 14283643262 | 38348422 | 37812563 | 535859 | 1.40 | 5.7 | 5.6 | 46 | 36 |
| RP121_1 | heat | D5 | 13625006123 | 36580213 | 36043152 | 537061 | 1.47 | 5.4 | 5.3 | 44 | 36 |
| RP121_2 | heat | D5 | 13625006123 | 36580213 | 36043152 | 537061 | 1.47 | 5.4 | 5.3 | 44 | 36 |
| RP122_1 | heat | D3 | 16076158335 | 43161019 | 42625226 | 535793 | 1.24 | 6.4 | 6.3 | 45 | 36 |
| RP122_2 | heat | D3 | 16076158335 | 43161019 | 42625226 | 535793 | 1.24 | 6.4 | 6.3 | 45 | 36 |
| RP130_1 | heat | D1 | 13614565840 | 36552173 | 36044664 | 507509 | 1.39 | 5.4 | 5.3 | 46 | 36 |
| RP130_2 | heat | D1 | 13614565840 | 36552173 | 36044664 | 507509 | 1.39 | 5.4 | 5.3 | 46 | 36 |
| RP141_1 | control | D3 | 12326927737 | 33095161 | 32575078 | 520083 | 1.57 | 4.9 | 4.8 | 46 | 36 |
| RP141_2 | control | D3 | 12326927737 | 33095161 | 32575078 | 520083 | 1.57 | 4.9 | 4.8 | 46 | 36 |
| RP144_1 | control | D5 | 11949656862 | 32082287 | 31624594 | 457693 | 1.43 | 4.8 | 4.7 | 45 | 36 |
| RP144_2 | control | D5 | 11949656862 | 32082287 | 31624594 | 457693 | 1.43 | 4.8 | 4.7 | 45 | 36 |
| RP159_1 | control | D1 | 11311705486 | 30369485 | 29954078 | 415407 | 1.37 | 4.5 | 4.4 | 46 | 36 |
| RP159_1 | control | D1 | 11311705486 | 30369485 | 29954078 | 415407 | 1.37 | 4.5 | 4.4 | 46 | 36 |

**Table S7. MIQE checklist for RT-qPCR.** Plants for RT-qPCR were grown alongside plants for RNA-seq analysis. Leaf discs were collected from 3 independent biological replicates per treatment per timepoint (days 1, 3 and 5 of heatwave) similarly to RNA-seq sampling (Methods). Detailed information for RNA extraction and RT-qPCR analysis are listed below in accordance with the MIQE guidelines. For primer sequences and information seeTable S8.

| **MIQE checklist** (as per Bustin *et al.* (2009)) | |
|---|---|
| **Experimental design** | |
| Definition of experimental and control groups | Experimental group: cowpea plants exposed to heat treatment<br>Control group: cowpea plants grown under control conditions |
| Number within group | 3 independent biological replicates per group, each grown in different cabinets. Each cabinet contained a separate plant for each timepoint. 3 technical replicates were performed for RT-qPCR. |
| **Sample** | |
| Description | Cowpea leaf material |
| Processing | 0.55 $cm^2$ cork borer used to cut leaf discs, immediately snap frozen in liquid nitrogen, stored at -80 °C for <6 months before RNA extraction |
| **Nucleic acid extraction** | |
| Procedure | A pestle and mortar were pre-cooled by adding liquid nitrogen. Once nearly all evaporated, the leaf disc sample was added to the mortar and ground to a fine powder. 20-30 mg were then used for RNA extraction. |
| Kit | NucleoSpin™ RNA Plant Kit (Macherey-Nagel) |
| DNase treatment | On-column treatment included as part of the kit (above). Each column was treated with 95 µl DNase solution for 15 min at room temperature. |
| RNA assessment | Purity and yield were measured using an LVis plate with SpectroStar Nano microplate reader (BMG Labtech) by evaluating absorbance ratios at 260/280 and 260/230 nm. Extractions with yield > 140 ng/µl, a 260/280 ratio near 2.0 and a 260/230 ratio >1.8 were selected. |
| **Reverse transcription** | |
| Complete reaction conditions | A subsample of 1 µg RNA was added to a 10 µl reaction containing 0.5 µl oligo-dT and 0.5 µl random nonamer primers, incubated for 5 min at 65 °C and immediately cooled on ice. Buffer, dNTP's, nuclease-free water and nanoScript2™ enzyme were added according to the instructions in the Precision nanoScript™ Reverse Transcription Kit (Primer Design). The final 20 µl reaction was incubated for 20 min at 42 °C, then 10 min at 75 °C. All cDNA was stored at -20 °C and diluted 1:5 prior to running RT-qPCR. |
| **RT-qPCR target and oligonucleotide information** | |
| See Supplementary Table 4 for gene IDs, primer sequences and amplicon lengths.<br>Primers were manufactured by Integrated DNA Technologies™ (IDT) and purified by desalting. | |
| **RT-qPCR protocol** | |
| RT-qPCR conditions | Hot start 95 °C for 2 min, then 40 cycles at 95 °C for 15 s and 60 °C for 1 min. |
| Melt curve | 95 °C for 1 min, 60 °C for 30 s and 95 °C for 30 s |
| Reaction volume | 15 µl |

| | |
|---|---|
| cDNA amount | 4 µl (40ng) |
| Master mix | PrecisionPLUS qPCR Master Mix (Primer Design) |
| Primer concentration | 0.467 µM |
| RT-qPCR instrument | AriaMx Real-Time PCR System (Agilent) |
| **RT-qPCR validation** | |
| Specificity | Checked for single peak in melt curve.<br>PCR products run on a gel and sequenced. |
| Primer efficiency / slope / y-intercept / $R^2$ of linear regression of $C_q$ versus ln(cDNA) / $C_q$ for NTC / $C_q$ for minus RT control | Primer efficiencies were calculated as described by Pfaffl (2001).<br>$C_q$ threshold = 450<br>NA where $C_q$ did not reach threshold |
| *HSP20* | 2.06 / -3.19 / 17.97 / 0.9970 / NA / 37.60 |
| *Rca1β* | 1.98 / -3.37 / 13.95 / 0.9993 / NA / 28.25 |
| *Rca8α* | 1.97 / -3.39 / 16.77 / 0.9965 / NA / NA |
| *Rca10α* | 1.91 / -3.57 / 18.61 / 0.9991 / NA / 30.33 |
| *Rca10β* | 2.03 / -3.25 / 23.95 / 0.9971 / NA / 28.47 |
| *Pp2A* | 2.01 / -3.30 / 21.75 / 0.9989 / NA / 27.17 |
| *Ubq28* | 1.95 / -3.44 / 19.80 / 0.9985 / NA / NA |
| *PolyP* | 1.85 / -3.74 / 23.70 / 0.9992 / NA / NA |
| *Elf1A* | 2.02 / -3.27 / 18.56 / 0.9934 / 37.67 / 31.83 |
| *B-Actin* | 1.95 / -3.45 / 23.22 / 0.9981 / NA / 30.30 |
| *Tua4* | 2.04 / -3.24 / 22.95 / 0.9963 / NA / 28.67 |
| **Data analysis** | |
| Analysis software | AriaMx, version 2.1 |
| Normalisation | The normalized relative quantity (NRQ) of expression was calculated in relation to the quantification cycle ($C_q$) values and the primer efficiency (E) of the target gene (goi) and the reference genes (ref1, ref2, ref3), based on Rieu & Powers (2009): $NRQ = \frac{E_{goi}^{-Cq}}{\sqrt{E_{ref1}^{-Cq} \cdot E_{ref2}^{-Cq} \cdot E_{ref3}^{-Cq}}}$ |
| Number and justification of choice of reference genes | Six cowpea reference genes were selected from the literature (Da Silva *et al.*, 2015; Weiss *et al.*, 2018) and assessed for stability across 10 experimental samples that varied in terms of treatment (control and heat-treated samples) and leaf age. Both geNorm (Vandesompele *et al.*, 2002) (qbase+, Biogazelle) and Normfinder (Andersen *et al.*, 2004) software were used to analyse the results. The three most stable genes, Pp2a, PolyP and Ubi28, were used to normalise gene expression. |
| Statistical method | NRQ values were log transformed and evaluated for statistical significance using Two-Way ANOVA followed by post-hoc Tukey's test. |

216 **Table S8. RT-qPCR primers.**

| Type | Gene/transcript ID | Primer Name | Sequence (5’ > 3’) | Amplicon length (bp) |
|---|---|---|---|---|
| Control | Vigun06g055220 | Vu06g052200_HSP20_F | GCTTCACCTAAAGTGCTGTTGA | 147 |
| | | Vu06g052200_HSP20_R | CTTACTGTATTCATCGTCTCCTGC | |
| GOI | Vigun01g219300 | VuRca1.1_F_v1 | CTTGGAAATGCTAACGAAGATGC | 156 |
| | | VuRca1.1_R_v2 | CACAAGTGCCATCACAATTGC | |
| GOI | Vigun08g150700 | VuRca8.1_F_v5 | CTACCAGAGACGACCGAATTGG | 128 |
| | | VuRca8.1_R_v5 | CCTGAGTGCACCAAAGAAATCA | |
| GOI | Vigun10g051600.1 | VuRca10.1_F_v2 | GAAACTTCTATGGACAAGCAGCT | 134 |
| | | VuRca10.1_R_v2 | CATCACCTAAAGTGTGTATGTGCA | |
| GOI | Vigun10g051600.2 | VuRca10.2_F_v1 | TGCTTGTCCAAGAGCAAGAGA | 144 |
| | | VuRca10.2_R_v1 | ACAACTTTAGGATCAAATCAAGGGT | |
| Reference | Vigun07g271800 | VuPp2A_F_v2 | TCAGCATATTCTTCCTTGTGTGA | 101 |
| | | VuPp2A_R_v2 | CTAAGACTGGTGCCATTCCC | |
| Reference | Vigun06g121000 | VuUbq28_F | GAGCTCAAGGACCTCCAGAA | 130 |
| | | VuUbq28_F | CTAGAAAAACACCCCCAGCA | |
| Reference | Vigun06g003800 | VuPolyP_F | CATGCAGACCACAAGGATTGA | 172 |
| | | VuPolyP_R | GAGGAGGTGACTGGCACAT | |
| Reference | Vigun04g088900 | VuElf1A_Fv_2 | GCTGTAACAAAATGGATGCCAC | 120 |
| | | VuElf1A_Rv_2 | CAAATGGAATCTTGTCTGGGTTG | |
| Reference | Vigun04g203000 | VuBAct_F_v2 | CTTCCAGCAGATGTGGATTGC | 146 |
| | | VuBAct_R_v2 | GCAGGCAGCAGTTGTTCC | |
| Reference | Vigun03g291000 | VuTua4_F_v2 | GGCGTTCCCTTGACATTGAG | 182 |
| | | VuTua4_R_v2 | GGGAGCATAAGATGAAAGCATAAAG | |

217

**Table S9. Modelling of the temperature response of Rubisco activation by cowpea Rca in leaf extracts of control and heat-treat plants.** The model providing the best fit to the data is highlighted in bold and was selected according to the lowest AIC score (Akaike information criterion) calculated according to using the AIC function in R. Models were applied to the full dataset shown in Fig. 4 (n=6-8).

| Treatment | Rubisco activation | | |
|---|---|---|---|
| | Model | degrees of freedom (K) | AIC score |
| Control | $2^{nd}$ order poly | 4 | 246.3 |
| | $3^{rd}$ order poly | 5 | 221.0 |
| | **$4^{th}$ order poly** | **6** | **216.9** |
| | $5^{th}$ order poly | 7 | 218.6 |
| | GAM | 3 | 280.6 |
| Heat | $2^{nd}$ order poly | 4 | 235.3 |
| | $3^{rd}$ order poly | 5 | 187.3 |
| | **$4^{th}$ order poly** | **6** | **186.5** |
| | $5^{th}$ order poly | 7 | 187.35 |
| | GAM | 3 | 278.6 |

**Table S10. Leaf total soluble protein (TSP) and chlorophyll content.** Leaf extracts used were aliquoted from the Rca temperature response measurements. TSP (n= 55-56), total chlorophyll (n= 54), chlorophyll a/b (n=50-55). All analysis conducted on the fifth day of heatwave. p-value and t-value determined from t-test (ns: $p > 0.05$, *: $p \leq 0.05$, **: $p \leq 0.01$, ***: $p \leq 0.001$).

| Data Set | Treatment | Mean | p-value | t-value |
|---|---|---|---|---|
| TSP (µg/µl) | control | 3.65 ± 0.45 | 0.002 (**) | 3.163 |
| | heat | 3.35 ± 0.53 | | |
| Total chlorophyll (mg/m$^2$) | control | 434 ± 42.3 | 0.455 (ns) | -0.750 |
| | heat | 439 ± 35.1 | | |
| Chlorophyll a/b | control | 2.22 ± 0.04 | 0.000183 (***) | -3.886 |
| | heat | 2.26 ± 0.06 | | |
